# Genetic architecture and genomic prediction accuracy of apple quantitative traits across environments

**DOI:** 10.1101/2021.11.29.470309

**Authors:** Michaela Jung, Beat Keller, Morgane Roth, Maria José Aranzana, Annemarie Auwerkerken, Walter Guerra, Mehdi Al-Rifaï, Mariusz Lewandowski, Nadia Sanin, Marijn Rymenants, Frédérique Didelot, Christian Dujak, Carolina Font i Forcada, Andrea Knauf, François Laurens, Bruno Studer, Hélène Muranty, Andrea Patocchi

**Author notes:** **Email addresses (in order of authors)** ‘ ’; ‘ ’; ‘ ’; ‘ ’; ‘ ’; ‘ ’; ‘ ’; ‘ ’; ‘ ’; ‘ ’; ‘ ’; ‘ ’; ‘ ’; ‘ ’; ‘ ’; ‘ ’; ‘ ’.

## Abstract

Implementation of genomic tools is desirable to increase the efficiency of apple breeding. The apple reference population (apple REFPOP) proved useful for rediscovering loci, estimating genomic prediction accuracy, and studying genotype by environment interactions (G×E). Here we show contrasting genetic architecture and genomic prediction accuracies for 30 quantitative traits across up to six European locations using the apple REFPOP. A total of 59 stable and 277 location-specific associations were found using GWAS, 69.2% of which are novel when compared with 41 reviewed publications. Average genomic prediction accuracies of 0.18–0.88 were estimated using single-environment univariate, single-environment multivariate, multi-environment univariate, and multi-environment multivariate models. The G×E accounted for up to 24% of the phenotypic variability. This most comprehensive genomic study in apple in terms of trait-environment combinations provided knowledge of trait biology and prediction models that can be readily applied for marker-assisted or genomic selection, thus facilitating increased breeding efficiency.

## Introduction

Apple (*Malus domestica* Borkh.) is the third most produced fruit crop worldwide^1^. Since its domestication in the Tian Shan mountains of Central Asia, the cultivated apple developed into a separated near-panmictic species^2^. Over the centuries, thousands of apple cultivars have been raised and conserved thanks to grafting^3^. Extensive relatedness among cultivars with a strong influence of a few founders through the history of apple breeding has been reported despite their high genetic diversity^4–6^. Only a fraction of the existing cultivars are grown commercialy^3^ and they require an intensive use of pesticides for crop protection. To diversify apple production, it is desirable to produce new cultivars for sustainable intensive agriculture and adapted to future climate, while remaining attractive to consumers.

Apple breeding is labor- and time-intensive, but selection efficiency can be improved by integrating DNA-informed techniques into the breeding process^7^. Marker-assisted selection allows breeders to predict the value of a target trait based on its association with a genetic marker. The method leads to removal of inferior seedlings without phenotyping, thus reducing the labor costs when decreasing the number of individuals passing to the next selection step^7^. Quantitative trait locus (QTL) mapping has been traditionally used to investigate the genetic basis of variation in traits such as pathogen resistance, phenology, and some fruit quality traits^8–11^. To bridge the gap between the discovery of marker-trait associations and their application in breeding, protocols that transfer the knowledge obtained by QTL analyses into DNA tests were established^12,13^. However, marker-assisted selection in apple remains restricted to a limited number of traits associated with single genes or a handful of large-effect QTL, such as pathogen resistance and fruit firmness, acidity, or color^14^. DNA-informed selection is rarely deployed in apple when breeding for quantitative traits with complex genetic architecture, though this task became feasible with the recent technological developments in apple genomics.

In the genomics era, advancements in genotyping and sequencing technologies led to a broad range of new tools for genetic analyses. In the case of apple, several reference genomes have been produced^15–19^, single nucleotide polymorphism (SNP) genotyping arrays of different densities such as 20K or 480K SNPs have been developed^20,21^, and genotyping-by-sequencing methods have been adopted^22,23^. Genome-wide association study (GWAS) emerged as a method for exploring the genetic basis of quantitative traits^24^. GWAS in apple have been used to identify associations between markers and various traits such as fruit quality and phenology traits^22,23,25-29^. The associations found with GWAS can be translated into DNA tests for marker-assisted selection. Besides GWAS, genomic selection was developed to exploit the effects of genome-wide variation at loci of both large and low effect on quantitative traits using a single model^30^ and is sometimes called marker-assisted selection on a genome-wide scale^31^. For genomic selection, prediction models are first trained with phenotypic and genomic data of a training population. In a second step, the models predict the performance of breeding material based on the genomic data alone. These genomic estimated breeding values are then used to make selections among the breeding material, thus increasing the breeding efficiency and genetic gain. Several studies have assessed genomic prediction accuracy for apple quantitative traits related to fruit quality and phenology^22,23,29,32-36^. Genomic selection can double genetic gain, as demonstrated by yield traits in dairy cattle^37^, but the accuracy of genomic prediction for yield traits in apple has not been studied. Analyses of genomic datasets beyond 100K SNPs have been limited to flowering and harvest time (GWAS and genomic prediction)^26,36^, fruit firmness and skin color (GWAS)^28,38^. Marker density, trait architecture, and heritability have been shown to differentially affect prediction performance in simulated data and for apple^34,36,39^ and their impact on genomic analyses should therefore be further empirically tested. Moreover, GWAS for the same traits measured at different locations, the effect of genotype by environment interaction (G×E) on genomic prediction accuracy, and predictions with multivariate genomic prediction models have not been evaluated yet in apple.

Plants are known for their strong phenotypic response to environmental factors, a phenomenon regularly tested in plant breeding using multi-environment trials. In general, when statistical models are applied to measurements from multi-environment trials, the effect of environment on individuals remains constant at single locations, but the G×E leads to changes in the ranking of genotypes across locations. With an increasing proportion of G×E effect relative to genotypic effect, both heritability and response to selection decrease^40^. A noticeable effect of contrasting European environments and G×E on two apple phenology traits – floral emergence and harvest date – has been reported, which demands testing the multi-environment modelling approaches in apple^36^. A location-specific GWAS may be used to identify loci with stable effects across environments and loci specific to individual locations^41^. Multi-environment prediction models can account for G×E by explicitly modeling interactions between all available markers and environments^42^. Borrowing information from other genotypes across environments through markers, the G×E method can outperform more simple modelling approaches that ignore G×E^42–44^. Additionally, taking advantage of information that traits provide about one another, a multivariate (also called multi-trait) genomic prediction can be applied. This method may be useful in case the assessment of one trait remains costly, but another correlated trait with less expensive measurement is available or can be assessed more easily^45^. The multivariate prediction can also be extended to a multi-environment approach when treating measurements from different environments as distinct traits^46^.

A population of 269 diverse apple accessions from across the globe and 265 progeny from 27 parental combinations originating in recent European breeding programs constitutes our apple reference population (apple REFPOP)^36^. The apple REFPOP has a high-density genomic dataset of 303K SNPs and was deemed suitable for the application of genomics-assisted breeding^36^. Combined with extensive phenotypic information, the apple REFPOP provides the groundwork for marker-assisted and genomic selection across contrasting European environments. Hence, 30 traits related to productivity, tree vigor, phenology, and fruit quality were measured in the apple REFPOP during up to three years and at up to six locations with various climatic conditions of Europe (Belgium, France, Italy, Poland, Spain, and Switzerland). First, GWAS was performed to dissect the genetic architecture of the studied traits, identify associated loci stable across locations and location-specific loci, and to observe signs of selection on loci of large effect. Second, this study aimed to measure prediction accuracy for these traits using single-environment univariate, single-environment multivariate, multi-environment univariate, and multi-environment multivariate genomic prediction models. Finally, a critical analysis of our results provided recommendations for future implementation of genomic prediction tools in apple breeding.

## Results

### Phenotypic data analysis

The accession and progeny groups of the apple REFPOP were evaluated for 30 quantitative traits at up to six locations. The measurements for ten traits were collected at one location, while the remaining 20 traits were available from at least two locations (three traits were measured in two locations, three traits in four locations, eleven traits in five locations and three traits in six locations, Supplementary Table 1). Most traits (25) were assessed during three seasons while five traits were measured during two seasons (Supplementary Table 1). Accounting for environmental effects in the phenotypic data, BLUPs of traits (best linear unbiased prediction of random effects of genotypes, see Equation 1) were produced across all locations and separately for each location. The traits showed unimodal as well as multimodal distributions (Supplementary Figure 1). Differences of various extent between the accession and progeny groups were observed (Supplementary Figure 2). As expected, high phenotypic and genotypic correlations (>0.7) between traits were observed within trait categories, namely the phenology, productivity, fruit size, outer fruit, inner fruit, and vigor category (Figure 1a). A few moderate positive phenotypic correlations (0.3–0.7) were found between trait categories such as harvest date and fruit firmness (0.51), yellow color and russet cover (0.55), soluble solids content and russet cover (0.36), or between yield (weight and number of fruits) and vigor trait category (0.36–0.51, Figure 1a). High average correlations were observed between the environments (combinations of location and year) for harvest date (0.82 [0.73, 0.95]) or red over color (0.80 [0.62, 0.92]) whereas low average correlations (<0.3) were present between environments for flowering intensity (0.18 [-0.49, 0.68]) and trunk increment (0.16 [-0.31, 0.55], Supplementary Table 2, Supplementary Figure 3). A shift of the progeny group compared to the accession group towards smaller, more numerous and less russeted fruits was observed (Figure 1b).

**Figure 1:**
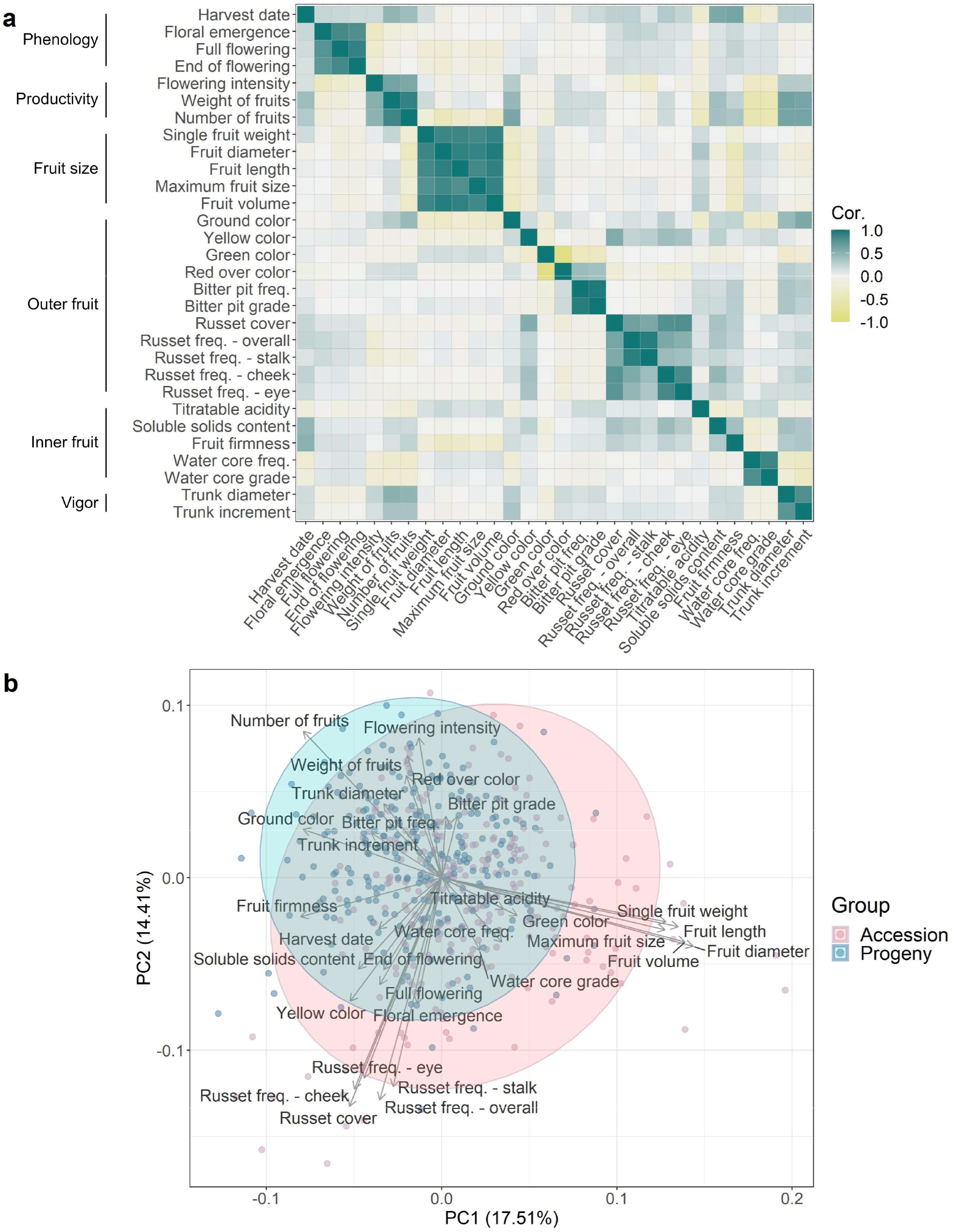
Exploratory phenotypic data analysis of the studied quantitative apple traits. **a** Pairwise correlations between traits with the phenotypic and genomic correlations in the lower and upper triangular part, respectively. Phenotypic correlation was assessed as Pearson correlation between pairs of across-location BLUPs, the genomic correlation as Pearson correlation between pairs of genomic BLUPs estimated from a G-BLUP model. Trait categories are outlined along the vertical axis. **b** Principal component analysis biplot based on across-location BLUPs of apple traits with the addition of location-specific BLUPs for traits measured at a single location.

### Genome-wide association studies

Across-location GWAS for 20 traits measured at more than one location (Supplementary Table 1) and location-specific GWAS for all 30 traits were used to explore the genetic basis of the assessed traits. The quantile-quantile plots showed that the observed and expected distributions of p-values corresponded well and no apparent inflation of p-values was found (Supplementary Figure 4 and 5). Across-location GWAS revealed 59 significant (-*log*_10_(*p*) > 6.74) marker-trait associations in 18 traits (Figure 2a, Supplementary Table 3). No significant associations were observed for trunk diameter and russet cover in the across-location GWAS. In the location-specific GWAS, 309 significant marker-trait associations for all 30 traits were discovered (Figure 2b, Supplementary Table 3). Of these 309 marker-trait associations, 32 associations for twelve traits were shared between the location-specific GWAS and the across-location GWAS (Supplementary Table 3). The coefficient of determination (*R*^2^) of significant associations was the largest for red over color (0.71), green color (0.55) and harvest date (0.42, Figure 2c, Supplementary Table 3).

**Figure 2:**
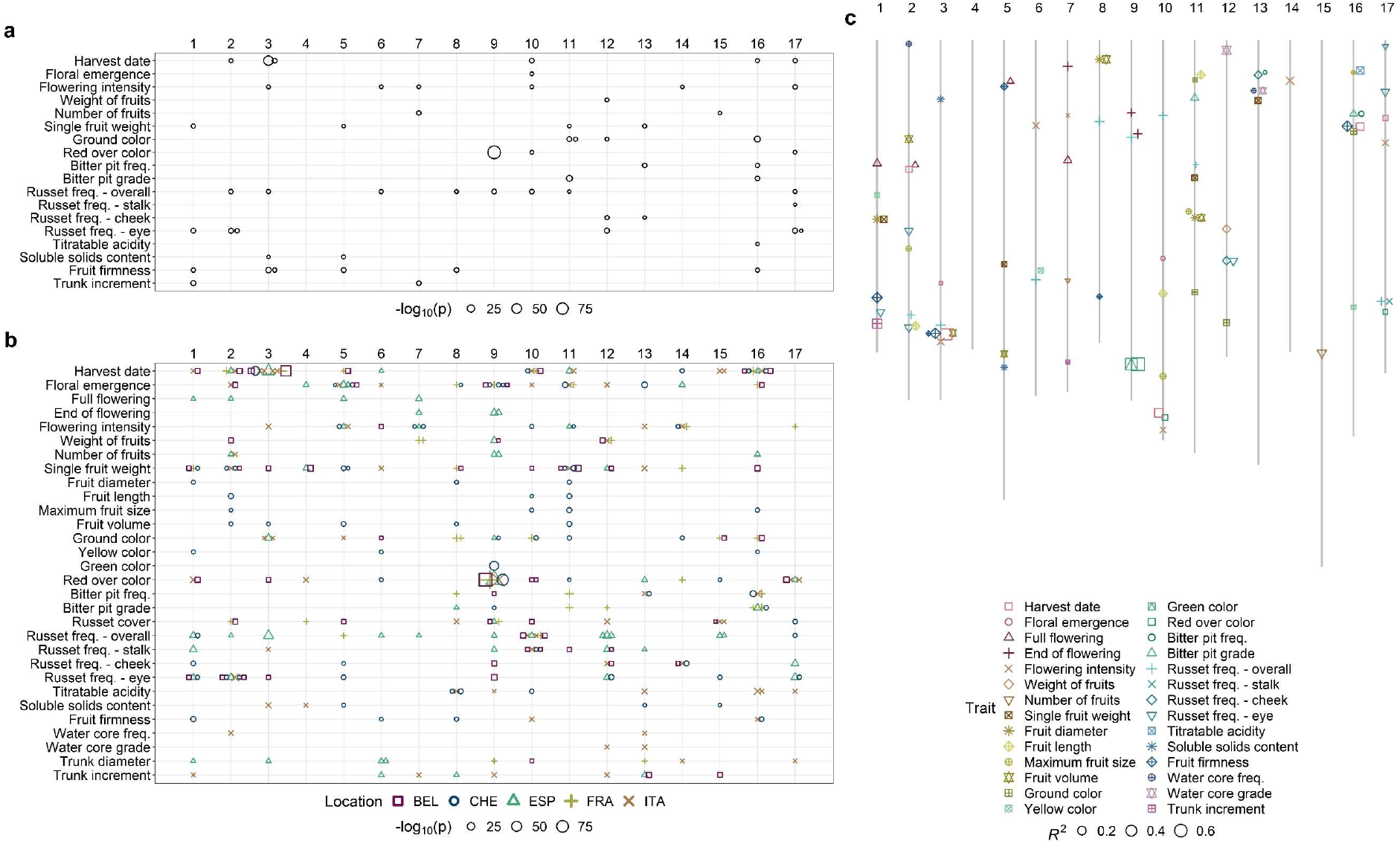
Significant marker-trait associations found by GWAS. **a** Distribution of the significant associations and corresponding p-values from across-location GWAS over the 17 apple chromosomes. **b** Distribution of the significant associations and corresponding p-values from location-specific GWAS over the 17 apple chromosomes. Locations are labeled as BEL (Belgium), CHE (Switzerland), ESP (Spain), FRA (France) and ITA (Italy). **a-b** Size of the symbols indicate the -*log*_10_(*p*). The x-axis shows chromosome numbers. **c** Physical positions (in bp) of the significant associations on chromosomes with their respective coefficients of determination (*R*^2^) from the across-location GWAS complemented with the location-specific GWAS for traits measured at a single location. Size of the symbols indicate the *R*^2^. The x-axis shows chromosome numbers.

Significant associations with different traits co-localized at identical positions or occurred very close in some genomic regions (distance between marker positions below 100 kb, Figure 2c, Supplementary Table 3). In the across-location GWAS, a marker significantly associated with harvest date on chromosome 3 (position 30,681,581 bp) was located next to two markers associated with fruit firmness (positions 30,587,378 and 30,590,166 bp). The same marker on the position 30,681,581 bp was also associated with harvest date, ground color, overall russet frequency and soluble solids content measured at several different locations (location-specific GWAS). Similarly, the association with harvest date on chromosome 16 (position 9,023,861 bp) was closely located to a marker associated with fruit firmness (position 8,985,888 bp) in the across-location GWAS. The traits related to bitter pit analyzed in the across-location GWAS, i.e., bitter pit frequency and grade, showed significant associations on chromosome 16, position 7,681,416 bp. Several associations with traits measuring fruit skin russet in the across-location GWAS co-localized on chromosome 12 (position 23,013,281 bp, russet frequency on cheek and in the eye) and 17 (position 27,249,890 bp, overall russet frequency and russet frequency at stalk). A marker at position 18,679,105 bp on chromosome 1 was associated with both single fruit weight from the across-location GWAS and fruit diameter from Switzerland (found with the location-specific GWAS). The association with marker at position 2,005,502 bp on chromosome 8 was shared between fruit diameter and fruit volume from Switzerland and single fruit weight from Belgium. On chromosome 11, fruit diameter, fruit volume and single fruit weight from Switzerland, as well as single fruit weight from Belgium, shared the association at position 18,521,895 bp. Additionally, position 3,622,193 bp on chromosome 11 was shared between the associations of fruit length and single fruit weight from Switzerland. For red over color and green color, the association with a marker on chromosome 9 (position 33,799,120 bp) occurred in across-location and four location-specific GWAS, while a close marker (position 33,801,013 bp, less than 2kb away) was associated in the two other location-specific GWAS. Additional significant marker-trait associations occurred in the same genomic regions among the location-specific GWAS and between the across-location and location-specific GWAS (Supplementary Table 3).

Previous reports on QTL mapping and GWAS in apple were extensively reviewed and 41 publications reporting on traits measured similarly to our own were found and taken for comparison (Supplementary Table 4). The QTL positions from literature and the marker-trait associations found in this study were assigned to chromosome segments (top, center, and bottom of a chromosome). Unique segment-trait combinations were discovered in the literature (166), in the across-location GWAS (52) and in the location-specific GWAS (172, Figure 3a). Out of all segment-trait combinations across our GWAS, 30.8% overlapped with the previously published results of QTL mapping or GWAS and the rest (69.2%) were novel. All previously published segment-trait combinations for the trait groups bitter pit and trunk were also detected in our study, whereas no overlap between the former and present associations was found for ground color and sugar trait groups (Figure 3b, Supplementary Figure 6).

**Figure 3:**
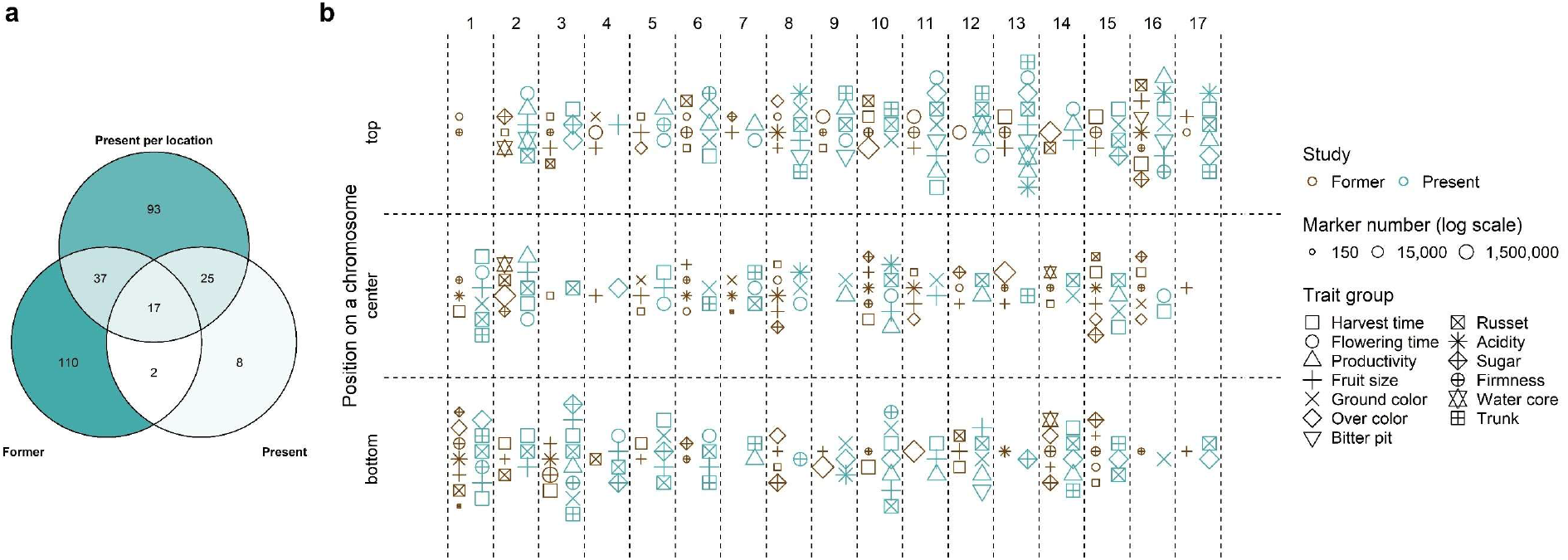
Comparison of the significant marker-trait associations with previously published associations. **a** Venn diagram comparing the unique associations, which were either previously published (former), reported in the across-location GWAS (present) or the location-specific GWAS (present per location). Color intensity and the values reflect the number of associations per diagram area. **b** Scatterplot of unique associations comparing published associations (former) with the merged across-location and location-specific GWAS (present). The traits were assembled into trait groups based on their similarity. Symbol size reflects the number of markers used in the studies. In case more than one publication reported an association in the same chromosome segment, only the report with the largest number of markers is shown (see Supplementary Table 4 for the complete list of previously published associations). **a-b** Positions of associations were assigned to three chromosome segments: top, center and bottom. Only the unique combinations of trait groups with segments and type of study (former or present) are shown.

### Allele frequency dynamics over generations

Eleven major significant marker-trait associations (*R*^2^>0.1) were identified in the global GWAS results (across-location GWAS with the addition of location-specific GWAS for traits measured at a single location only, Figure 4). Among these major associations, changes in the frequency of alleles with an increasing effect on trait phenotypes were quantified in 30 ancestral accessions (five ancestor generations of the progeny group, Supplementary Table 5) and all 265 progenies included in the apple REFPOP (Figure 4a). Compared to the ancestral accessions, the frequency of the allele with an increasing effect on phenotype (Figure 4c) was higher in the progeny for the alleles associated with later harvest date and increased flowering intensity, titratable acidity, fruit firmness and trunk increment (Figure 4a). For the marker associated with green color and red over color, the allele frequencies were equivalent for ancestors and progeny, which reflected the minor allele frequency of nearly 0.5 for both traits (Figure 4b,d). Noticeably, at the markers closely associated with harvest date and fruit firmness on chromosome 3, the allele associated with later harvest date and firmer fruits was fixed in all progeny, while the allele with a decreasing effect on the phenotype was present with a frequency below 0.1 in the whole apple REFPOP (Figure 4a-d). The allele associated with larger trunk increment on chromosome 1 was found in progeny known to segregate for *Rvi6*, and it was present in only two accessions (‘Prima’ and X6398) that are also known to carry the apple scab resistance gene *Rvi6*, which is located about 1.8 Mb from the SNP associated with trunk increment (Figure 4b-c). The remaining associations (*R*^2^≤0.1) reported by the global GWAS showed various trends in allele frequencies across generations such as increased frequency of alleles associated with increased weight of fruits in the progeny (Supplementary Figure 7). The individual parental combinations of the progeny group were often fixed for single alleles (Figure 4b, Supplementary Figure 8). Boxplots of the across-location BLUPs against the dosage of the reference allele (0, 1, 2) for the eleven major significant marker-trait associations showed additive effects of the alleles on phenotypes (Supplementary Figure 9). Squared Pearson’s correlations in a window of ~3,000 markers surrounding each of the major significant marker-trait associations showed that markers in linkage disequilibrium extended over larger distances around some marker-trait associations (Supplementary Figure 10). When visually compared with other loci, the associations with harvest date and fruit firmness on chromosome 3 as well as red over color and green color on chromosome 9 were found in genomic regions of the highest linkage disequilibrium between markers (Supplementary Figure 10). The markers associated with trunk increment and *Rvi6* also showed signs of linkage disequilibrium (Supplementary Figure 10).

**Figure 4:**
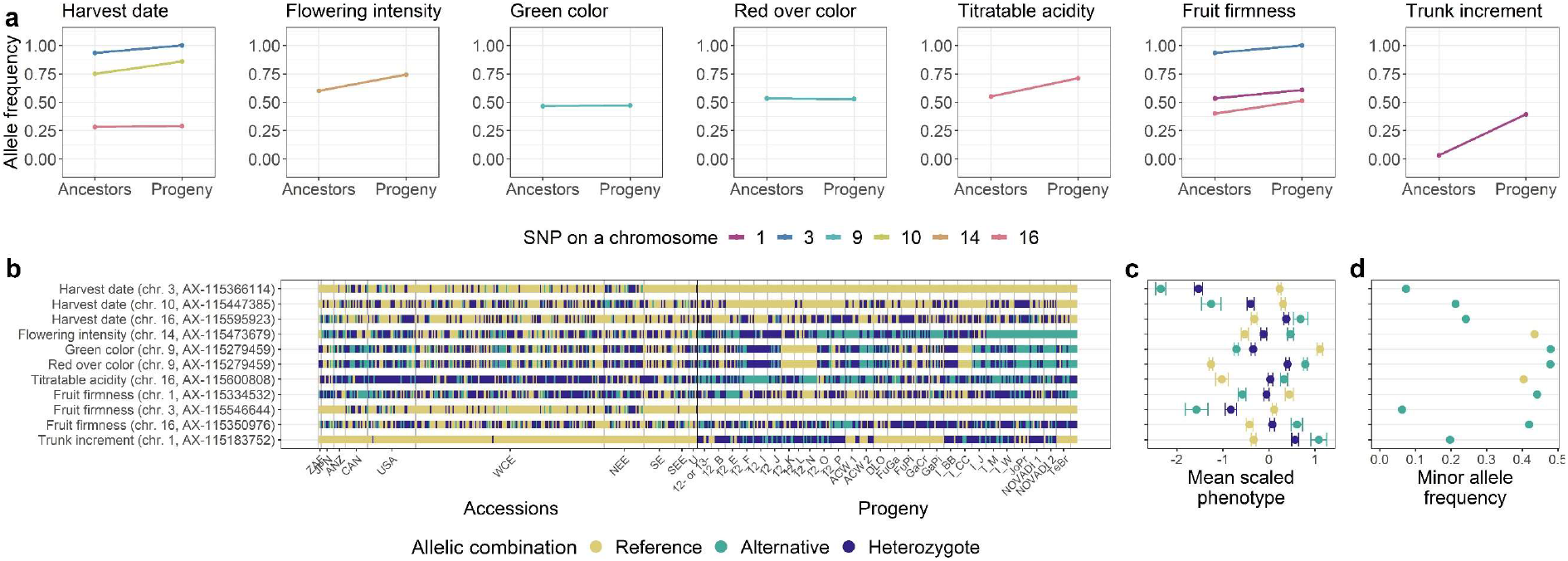
Allele frequency dynamics of the major significant marker-trait associations. **a-d** The associations were chosen based on the coefficient of determination (*R*^2^>0.1) from the global GWAS. **a** For each association, frequency of the allele with increasing effect on trait phenotypes in the apple REFPOP is shown. For the progeny group (progeny) and its five ancestor generations (ancestors), the allele frequencies are shown as points connected with a line. Out of all known ancestors, the allele frequency was estimated for 30 accessions included in the apple REFPOP. Colors of the points and lines correspond to chromosome locations of the associated SNPs. **b** Allelic combinations carried by the apple REFPOP genotypes, sorted according to geographic origin of accessions and affiliation of progeny to parental combinations (the x-axis was labeled according to Supplementary Table 1 and 2 in Jung et al.^36^). **c** Phenotypic BLUPs of traits and their standard error for each allelic combination, centered to mean 0 and scaled to standard deviation of 1. **d** Frequency of the minor allele in the whole apple REFPOP. **b-d** The legend and y-axis are shared between plots. In d, the color of an allele corresponds to the color of the homozygous allelic combination of the same allele in b and c.

### Genomic prediction

Four single-environment univariate prediction models – random forest (RF), BayesCπ, Bayesian reproducing kernel Hilbert spaces regression (RKHS) and genomic-BLUP (G-BLUP) – and a single-environment multivariate model with an unstructured covariance matrix of the random marker effect (MTM.UN) were compared using across-location BLUPs and location-specific BLUPs as phenotypes from a single environment. Among these models, the average prediction accuracies per trait 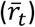 ranged between 0.18 for russet cover and 0.88 for red over color, both extreme values observed with RF (Supplementary Table 6). The prediction accuracies estimated for G-BLUP were further used as reference for model comparisons due to its widespread use in genomic prediction. When the prediction accuracy of the G-BLUP model was averaged over all traits 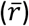, the obtained 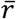 was equal to 0.50. The RF showed an 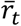 higher than G-BLUP for 9 out of 30 traits and an 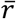 of 0.49. BayesCπ, RKHS and MTM.UN showed an 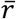 of 0.50, 0.51 and 0.50 and exceeded 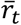 of G-BLUP in one, twelve and ten traits, respectively. Generally, a similar performance of all five models was observed (Figure 5a).

**Figure 5:**
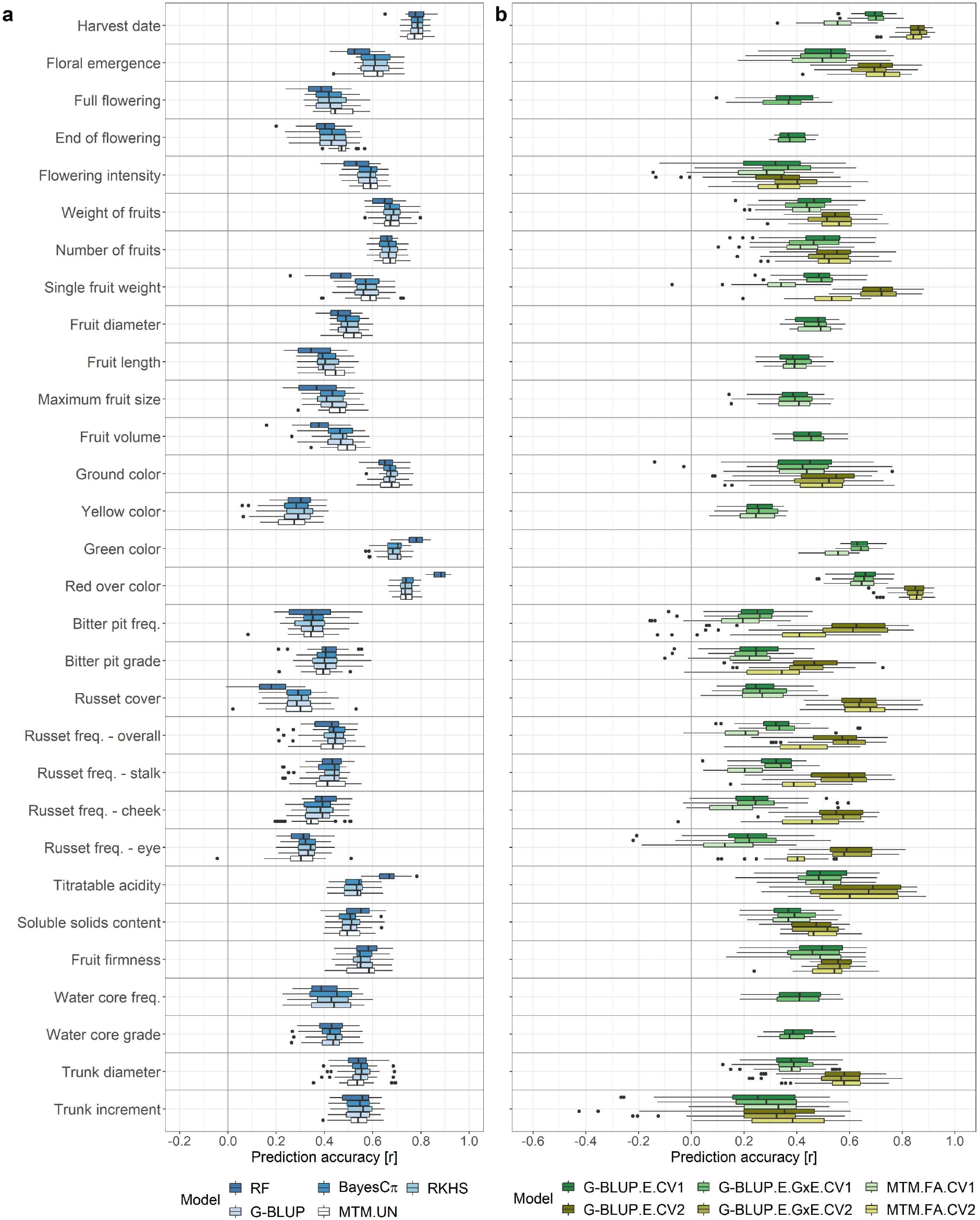
Genomic prediction accuracy in apple quantitative traits using eight genomic prediction models and two cross-validation scenarios. **a** Prediction accuracy of four single-environment univariate models, i.e., random forest (RF), BayesCπ, Bayesian reproducing kernel Hilbert spaces regression (RKHS) and genomic-BLUP (G-BLUP), and one single-environment multivariate model with an unstructured covariance matrix of the random marker effect (MTM.UN). The models were applied with a five-fold cross-validation where 20% of the genotypes were masked in each of the five runs. The MTM.UN was used in case a trait showed genomic correlation larger than 0.3 with at least one other trait. **b** Prediction accuracy of two multi-environment univariate models, i.e., across-environment G-BLUP (G-BLUP.E) and marker by environment interaction G-BLUP (G-BLUP.E.G×E), and the multi-environment multivariate factor-analytic model (MTM.FA). The models were applied under two five-fold cross-validation scenarios CV1 and CV2. The CV1 was applied for all traits using G-BLUP.E and G-BLUP.E.G×E and for traits measured in at least three environments using MTM.FA. The CV2 was applied for traits measured in Switzerland and in at least a one other location. **a-b** Prediction accuracy was estimated as a Pearson correlation coefficient between the observed and the predicted values of genotypes whose phenotypes were masked in a five-fold cross-validation. For the multi-environment models, the correlation coefficients were estimated for each environment separately. In the box plot, the bottom and top line of the boxes indicate the 25th percentile and 75th percentile quartiles (the interquartile range), the center line indicates the median (50th percentile). The whiskers extend from the bottom and top line up to 1.5-times the interquartile range. The points beyond the 1.5-times the interquartile range from the bottom and top line are labeled as dots.

When compared with the baseline model G-BLUP, the single-environment multivariate model MTM.UN showed an improved prediction accuracy for several traits when they were modelled in combination with a correlated trait (genomic correlation larger than 0.3, Figure 5a, Supplementary Table 6). The inclusion of floral emergence as correlated trait improved 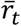 of full flowering and end of flowering. A combination with weight of fruits improved 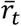 of flowering intensity. Fitting the model using fruit length showed an increased 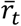 of single fruit weight and using single fruit weight led to an increase in 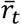 for fruit diameter, fruit length, maximum fruit size and fruit volume. Using soluble solids content resulted in an increase of 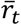 for russet cover, while using russet frequency at cheek led to an improved 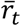 of russet frequency at stalk. Prediction accuracies for all possible combinations of correlated traits can be found in Supplementary Table 7.

Two multi-environment univariate models – across-environment G-BLUP (G-BLUP.E) and marker by environment interaction G-BLUP (G-BLUP.E.G×E) - and the multi-environment multivariate factor-analytic model (MTM.FA) were compared using two cross-validation scenarios corresponding to different experimental scenarios. In the first cross-validation scenario (CV1), traits were predicted for 20% of genotypes in each environment (i.e., their phenotypes were masked in all environments for model training). In the second cross-validation scenario (CV2), traits were predicted for 20% of genotypes in all but the Swiss environments (i.e., for these genotypes the environments “CHE.2018”, “CHE.2019” and “CHE.2020” were retained for model training). For the models applied with CV1, the 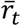 ranged between 0.13 (for russet frequency in the eye obtained with MTM.FA) and 0.70 (for harvest date estimated with G-BLUP.E.G×E, Supplementary Table 6). With CV2, the lowest 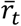 of 0.29 was measured for trunk increment with G-BLUP.E.G×E and the maximum 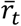 of 0.86 was found for harvest date with both G-BLUP.E and G-BLUP.E.G×E models (Supplementary Table 6). The prediction performance of G-BLUP.E, G-BLUP.E.G×E and MTM.FA was generally lower under CV1 than under CV2 (Figure 5b, Supplementary Table 6). For all traits, the G-BLUP.E.CV1, G-BLUP.E.G×E.CV1 and MTM.FA.CV1 showed lower 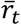 than the single-environment G-BLUP, the 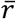 being equal to 0.40, 0.40 and 0.36, respectively. The G-BLUP.E.G×E.CV1 performed better than G-BLUP.E.CV1 for 14 out of 30 traits. The G-BLUP.E.CV2 and G-BLUP.E.G×E.CV2 outperformed G-BLUP for 13 out of 20 traits. The G-BLUP.E.CV2 and G-BLUP.E.G×E.CV2 both showed 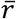 equal to 0.57. The increase in 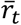 from G-BLUP to G-BLUP.E.CV2 (0.35) as well as from G-BLUP to G-BLUP.E.G×E.CV2 (0.36) was the most pronounced for russet cover. The performance of G-BLUP.E.CV2 and G-BLUP.E.G×E.CV2 remained below the level of G-BLUP predictions for productivity traits (flowering intensity, weight and number of fruits), ground color, soluble solids content, fruit firmness and trunk increment. The G-BLUP.E.G×E.CV2 performed better than G-BLUP.E.CV2 for 8 out of 20 traits. The 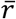 of MTM.FA.CV2 was equal to 0.52 and therefore similar to G-BLUP, however, the model outperformed G-BLUP for nine out of 20 predicted traits (Supplementary Table 6). The MTM.FA showed higher prediction accuracy than both G-BLUP.E and G-BLUP.E.G×E for two traits under CV1 and five traits under CV2 (Supplementary Table 6).

Across all model groups, the best prediction performance was found for harvest date, green color and red over color (Figure 5, Supplementary Table 6). The lowest prediction accuracy was found for traits related to bitter pit and russet as well as yellow color. Additionally, the prediction accuracy for flowering intensity and trunk increment with the multi-environment models remained strongly below the 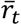 of the corresponding single-environment models.

### Synthesis of phenotypic and genomic analyses

The across-environment clonal mean heritability was generally very high in the evaluated traits, the value being close to one for harvest date and red over color and not lower than 0.80 for all the other traits with the exception of full flowering (0.74), end of flowering (0.79) and water core grade (0.79, Figure 6, Supplementary Table 6). The genomic heritability, which is the proportion of phenotypic variance explained by the markers, was larger than 0.80 for harvest date, floral emergence, green color and red over color, the value was not lower than 0.40 for all the other traits with the exception of bitter bit frequency (0.33) and grade (0.39, Figure 6, Supplementary Table 6).

**Figure 6:**
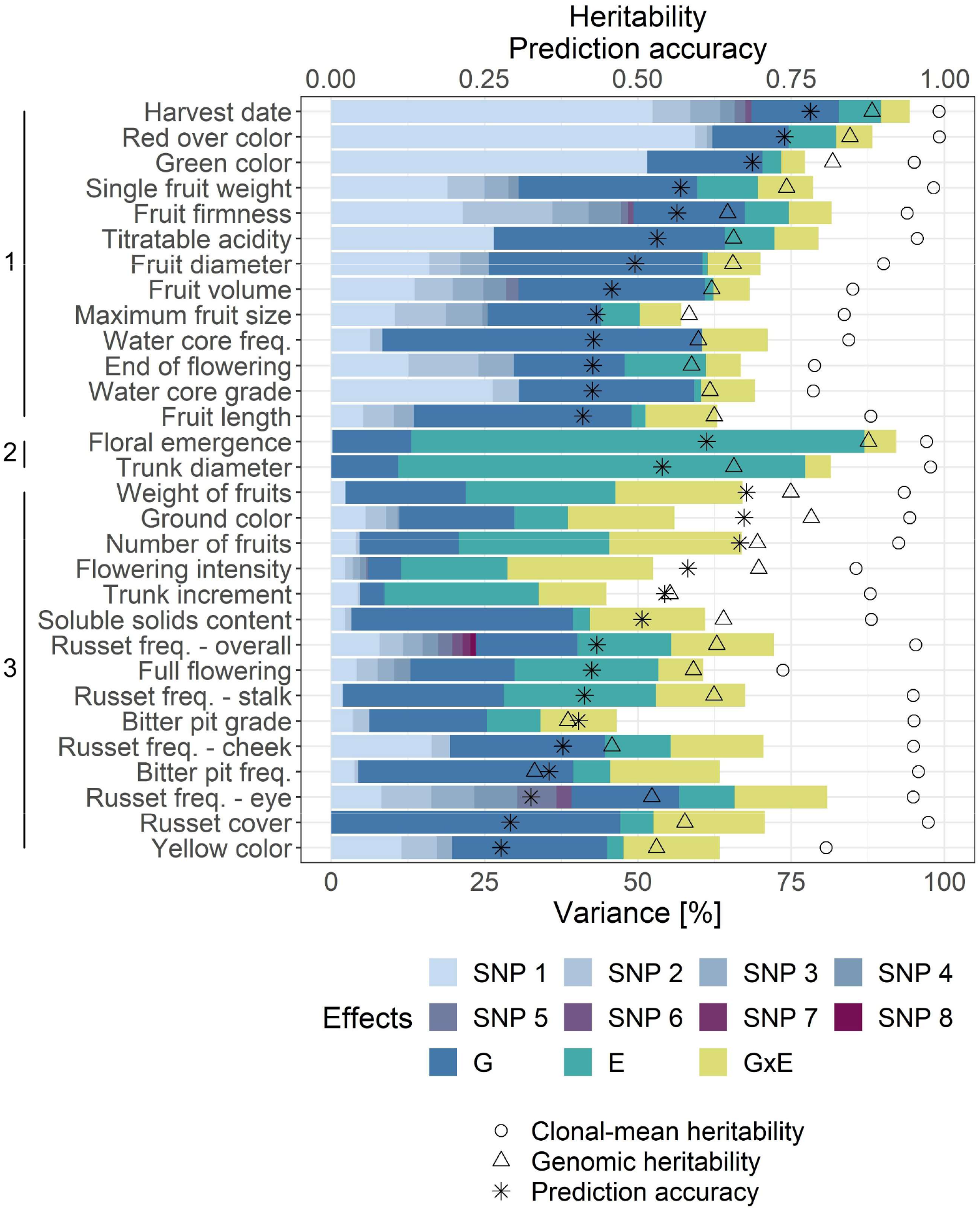
Synthesis of phenotypic and genomic analyses. Across-environment clonal mean heritability, genomic heritability, average prediction accuracy 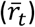 for the single-environment G-BLUP and the proportion of phenotypic variance explained by the effect of each significantly associated marker (SNP 1–8), genotype (G), environment (E) and genotype by environment interaction (G×E). The significantly associated markers corresponded to results of the global GWAS. Phenotypic variance components were used to estimate clusters of traits outlined along the vertical axis. Within each cluster, the traits were sorted according to 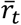.

The effects of genotype and significantly associated markers together explained a substantial part of the phenotypic variance of traits, the largest sums of these genotypic effects were observed for harvest date (82.8%) and red over color (74.6%, Figure 6, Supplementary Table 6). Altogether, the sum of the genotypic effects explained a very low proportion of the total variance for floral emergence (13.1%), flowering intensity (11.4%), trunk diameter (10.9%) and trunk increment (8.7%). The major proportion of the phenotypic variance was explained by the effect of environment for floral emergence (73.9%) and trunk diameter (66.3%). The lowest impact of environment was found for traits measured at only one location over two or three years such as fruit diameter or water core frequency, both showing an effect of environment (i.e., year) below 1%. The effect of G×E was the most pronounced for productivity traits, i.e., flowering intensity (23.7%), weight of fruits (20.8%) and number of fruits (21.6%).‮ The proportion of the G×E effect was the lowest for harvest date (4.7%), floral emergence (5.2%), red over color (5.9%) and trunk diameter (4.2%) among the traits measured at more than one location and for end of flowering (5.7%), fruit volume (5.9%) and green color (3.9%) among the traits measured at one location. A high proportion of the phenotypic variance remained unexplained by the model parameters for flowering intensity (47.5%), bitter pit grade (53.4%) and trunk increment (55.1%).

Hierarchical clustering of the phenotypic variance components revealed three clusters of traits (Figure 6). A strong genotypic effect and a comparably low effect of environment and G×E was observed for 13 traits assigned to the cluster one. Most of the phenotypic variance was explained by the effect of environment in floral emergence and trunk diameter, which were grouped in cluster two. Finally, 15 traits with a pronounced effect of environment and/or G×E were grouped in cluster three.

## Discussion

### Discovered loci overlap between association studies and traits

Our GWAS permitted to enlighten the architecture of analyzed traits as well as the identification of numerous marker-trait associations stable across, and specific to, the locations of the apple REFPOP. The particular design of the experiment, including the diversity of the plant material used (accessions and small progeny groups), multiple locations, and multiple years of evaluation, resulted in about two thirds of the discovered associations being novel when compared with the loci published in studies spanning more than two decades. Our study design also allowed us to replicate the identification of many previously known loci associated with the studied traits.

The association of one locus with two or more seemingly independent traits (i.e., caused by pleiotropy) and linkage disequilibrium between loci associated with different traits are frequent for complex traits^47^. The GWAS performed in this study showed several marker-trait associations at identical or close positions for different traits. The interdependency between harvest date and fruit firmness, which can be also observed empirically for early cultivars that soften more, may be an example of pleiotropy or linkage disequilibrium between loci. Harvest date and fruit firmness are known to be regulated by ethylene production^48^ and associated with loci present on chromosomes 3 (*NAC18.1*), 10 (*Md-ACO1, Md-PG1*), 15 (*Md-ACS1*) and 16^22-49–52^.

In this work, closely located (distance <100 kb) associations with both harvest date and fruit firmness were found on chromosome 3. Migicovsky et al.^22^ reported an overlap between associations with harvest time and fruit firmness on chromosome 3 falling within the coding region of *NAC18.1*. The authors hypothesized that the lack of associations on other chromosomes was likely due to low SNP density around the causal loci (the study used a GBS-derived 8K SNP dataset). The larger number of associations reported here might be a result of the high SNP density (303K SNPs) deployed in GWAS, however, not all previously reported loci were re-discovered.

The SNPs associated with harvest date and fruit firmness on chromosome 10 were further apart (~6 Mb). For harvest date, one of the associations on chromosome 10 was stable across locations and several associations were location specific. However, the association on chromosome 10 with fruit firmness was found for the Italian location only. It has been shown that chromosome 10 contains more than one QTL controlling fruit firmness^49–51^, but stable across-location association with fruit firmness on chromosome 10 was missing in our study. One of the known loci on chromosome 10, the *Md-PG1* gene, is responsible for the loss of fruit firmness after storage^51,53^. In apple REFPOP, fruit firmness was measured within one week after the harvest date and this very short storage period might have contributed to the less pronounced effect of the locus *Md-PG1* in our GWAS.

Two associations with harvest date measured in Italy but no association with fruit firmness were found on chromosome 15. Although a marker for *Md-ACS1* related to ethylene production was previously mapped on chromosome 15^50^, and QTL for fruit firmness was discovered on the same chromosome^49^, these markers did not co-locate, but rather, mapped at the opposite extremes of chromosome 15^49,50^. Likewise, the connection between harvest date and fruit firmness on chromosome 15 could not be confirmed here.

Our GWAS showed associations with harvest date and fruit firmness on chromosome 16, which were located 38 kb apart. In the past, loci associated with harvest date and fruit firmness have been reported in the same region on chromosome 16^26,49^. The role of this locus in the regulation of harvest date and fruit firmness remains unknown and requires further research.

In practice, ripeness of fruit (harvest date) is decided based on ground color and starch content. The GWAS results showed that the association on chromosome 3 was not only found for harvest date and nearby markers associated with fruit firmness, but also corresponded to associations with ground color and soluble solids content. This might be explained by the fact that these traits are used to define ripeness and thus harvest date. Further, the association of the *NAC18.1* locus on chromosome 3 with overall russet frequency would support the known enhanced expression of *NAC* transcription factors in russet skin^54^.

Co-localizations between associations found for different measures of bitter pit on chromosome 16, russet on chromosomes 12 and 17, fruit size on chromosomes 1, 8 and 11, and skin color on chromosome 9 are likely the result of relatedness among trait measurements. The measures that are easiest to score can be used in future to phenotype these traits.

### Signs of selection in marker-trait associations of large effect

The design of apple REFPOP allowed for the discovery of major marker-trait associations and for the analysis of changes in allele frequency between 30 ancestral accessions and 265 progeny included in the apple REFPOP. Comparing ancestors with the progeny, higher frequencies of the alleles associated with later harvest date and increased flowering intensity, titratable acidity, fruit firmness and trunk increment were found for the progeny. Of these traits, harvest date and fruit firmness are correlated, probably due to pleiotropy or linkage disequilibrium of causal loci, as it was shown in this and previous studies^22^. Consequently, the consistently higher frequency of alleles contributing to later harvest and firmer apples in the progeny is because the softening of harvested apples is undesirable and likely selected against^55^. Signs of selection for increased firmness were also recently found in USDA germplasm collection^5^. Our study also showed fixation of the late-harvest and high-firmness alleles on chromosome 3 in the whole progeny group, which suggests a loss of genetic diversity in the modern breeding material at this locus. For flowering intensity, a trait positively correlated with apple yield, a new locus was discovered on chromosome 14. The increased frequency of the allele contributing to higher flowering intensity in the progeny, its presence in all parental genotypes, and fixation in some parental combinations may be the result of breeding for high yield. The major locus found for acidity on chromosome 16 was consistent with the *Ma* locus frequently detected in various germplasm^8,11^. The total number of the high-acidity alleles for *Ma* and *Ma3*, which is another regularly detected acidity locus, was shown to be higher in parents of a European breeding program (Better3fruit, Belgium) than in parents used in the USDA breeding program^11,56^. The desired acidity level might depend on local climate of the breeding program and market preferences^56^. The increase in frequency of the allele contributing to higher acidity in the progeny may indicate a current preference towards more acidic apples in European breeding, but further investigation is needed to clarify the trend. The last locus of large effect showing allele frequency dynamics between generations was found for trunk increment. The allele associated with an increase in trunk increment may have been selected in the progeny due to its potential impact on productivity suggested by moderate positive correlations between tree vigor (trunk diameter and increment) and yield-related traits. Additionally, the marker associated with trunk increment was 1.8 Mb apart from a SNP marker associated with *Rvi6* gene responsible for resistance against apple scab^10^. These two markers (AX-115183752 for trunk increment and AX-115182989 (also called Rvi6_42M10SP6_R193) for apple scab) showed a correlation of 0.15 and occurred within a region of increased linkage disequilibrium between markers (Supplementary Figure 10). All accessions were homozygous for the reference allele of AX-115183752 associated with decreased trunk increment (Figure 6c) except for ‘Prima’ and X6398, which were heterozygous. The scab-resistant accessions ‘Prima’ and X6398 (which is a second-generation offspring of ‘Prima’^57^) but also ‘Priscilla-NL’ (known to be heterozygous for *Rvi6*^58^), were also heterozygous for AX-115182989. All other accessions were homozygous for the reference allele not associated with *Rvi6*. The allele on chromosome 1 associated with increased trunk increment may have been co-selected with the *Rvi6* locus responsible for resistance against apple scab.

Signs of intense selection for red skin were recently detected in the USDA germplasm collection when compared with progenitor species of the cultivated apple^5^. Our results show that the associations with red over color and green color, which phenotypically mirrored red over color and was associated with the same marker, did not show changes in allele frequency between ancestors and progeny included in the apple REFPOP. Some parental combinations showed almost exclusively the allele increasing red skin color, other parental combinations exhibited a lack of the allele. This uneven distribution of the alleles in the progeny group pointed to different directions of selection for fruit skin color in the European breeding programs (Figure 4b).

### Performance of the single-environment univariate genomic prediction models

Single-environment univariate genomic prediction models were applied to individual traits after accounting for environmental effects and averaging across locations and/or years. The observed small differences between genomic prediction accuracies of various models (Figure 5a) were in accordance with previous model comparisons where distinctions among models were negligible^39,59^. The largest extremes in prediction accuracy between traits were found with random forest, which allowed for the overall highest prediction accuracy among all compared models for red over color. The explanation for the striking performance of random forest for red over color might be found in the results of our GWAS. This trait of oligogenic architecture was associated with a few low-effect loci and one locus of large effect explaining 61% of the red over color phenotypic variance measured in the apple REFPOP. High correlations between many markers, i.e., linkage disequilibrium, were found in the vicinity of the large-effect locus (Supplementary Figure 10). Random forest is known to assign higher importance to correlated predictor variables (here the markers) in the tree building process^60^, which may have contributed to the particularly high prediction accuracy found for red over color with random forest.

The prediction accuracy for red over color reached ~0.4 in several former prediction studies^22,23,29,34^ and was approximately doubled in our work, which demonstrated the potential of the current study design for accurate genomic predictions. For harvest date, the currently reported prediction accuracy of 0.78 was only slightly higher than the accuracy of 0.75 obtained with the initial apple REFPOP dataset measured during one year^36^, but these accuracies showed a considerable improvement over other accuracies of approximately 0.5–0.6 reported elsewhere^22,23,29^. As shown before^36^, these results underline the suitability of apple REFPOP design for the application of genomic prediction.

Prediction accuracy for traits such as yellow color or russet cover were on the opposite side of the spectrum when compared to harvest date and red over color. The prediction accuracy of yellow color and russet cover was low, although the genotypic effects explained 45% and 47% of the phenotypic variance, respectively. The across-environment clonal-mean heritability of russet cover was high (0.97), while the heritability for yellow color was slightly lower (0.81, Figure 6). Yellow color showed a moderate phenotypic correlation of 0.55 with russet cover, suggesting that the phenotyping device might have classified some russet skin as yellow color. Symptoms of powdery mildew could have been misinterpreted as russet skin. The decreased performance of genomic prediction models might stem from inaccurate phenotyping methods, insufficient SNP density in the associated regions, or other factors, all of which could not be explained in this work.

### Role of genotype by environment interactions in multi-environment univariate genomic prediction

The multi-environment univariate genomic prediction models either directly estimated environmental effects (across-environment G-BLUP, called here G-BLUP.E) or additionally borrowed genotypic information across environments and thus considered the G×E (marker by environment interaction G-BLUP, called here G-BLUP.E.G×E)^42^. The average accuracy of the G-BLUP.E.G×E model across traits was only slightly higher than the accuracy of the G-BLUP.E. In contrast, the G-BLUP.E.G×E model had substantially greater prediction accuracy than the G-BLUP.E model when applied in wheat^42^. In the latter study, a productivity trait was measured under simulated conditions of mega-environments and the effect of G×E explained up to ~60% of the phenotypic variance^42^. Our work only focused on European environments and the largest proportion of phenotypic variance assigned to G×E was 24% for a productivity trait (flowering intensity). Furthermore, the average proportion of G×E across traits was approximately 12%, which may explain the mostly negligible differences between the G-BLUP.E and G-BLUP.E.G×E models. Our results were in line with the low interaction of additive genetic effects with location of up to ~6% obtained for apple fruit quality traits measured at two locations in New Zealand^33^, and the limited G×E reported for fruit maturity timing in sweet cherry across continents^61^. For approximately half of the tested traits, the G-BLUP.E.G×E did not outperform G-BLUP.E. For these traits, the G-BLUP.E ignoring G×E may be sufficient to account for the environmental effects across European sites because it is computationally simpler and therefore less demanding. Traits such as flowering intensity, soluble solids content, trunk increment or traits related to fruit size and russet showed an improved performance under G-BLUP.E.G×E when compared to G-BLUP.E. For traits positively responding to G-BLUP.E.G×E, the G×E should be considered when making predictions across environments. The highest improvement of prediction accuracy with G-BLUP.E.G×E when compared to G-BLUP.E was found for flowering intensity, the difference between the models amounting to 0.07 (Figure 5b). This result might be explained by the highest contribution of G×E to the phenotypic variance of flowering intensity among all traits (Figure 6). A comparably high contribution of G×E was also found for weight of fruits and number of fruits, though no improvement with G-BLUP.E.G×E model was observed for these traits. When comparing the relative contributions of variance components to the phenotypic variance of flowering intensity, weight of fruits and number of fruits, the genotype explained 11%, 22% and 21%, the environment 17%, 24% and 25%, and the G×E 24%, 21% and 22%, respectively. Although the proportions of G×E were similar in the three compared traits, the effects of genotype and environment explained a higher proportion of the variance for weight of fruits and number of fruits than for flowering intensity. This may have contributed to the surprisingly lower accuracy of the G-BLUP.E.G×E model when compared with G-BLUP.E for weight of fruits and number of fruits, but additional investigations may be needed to clarify this result in the future.

The G-BLUP.E.G×E model assumes positive correlations between environments and is therefore mostly suitable for the joint analysis of correlated environments^42,62^. As shown by Lopez-Cruz et al.^42^ and in our study, this assumption of G-BLUP.E.G×E resulted in the best model performance for traits showing high positive correlations between environments (here harvest date and red over color) and the worst performance for traits exhibiting low correlations between environments (here flowering intensity and trunk increment, Figure 5b, Supplementary Table 2, Supplementary Figure 3). For flowering intensity and trunk increment, bivariate prediction of the environments or prediction with a different G×E model not assuming positive correlations between environments might be more appropriate than the currently applied approach^42,63^.

### Multivariate models as a useful element in the genomic prediction toolbox

Multivariate (also called multi-trait) models were shown to be useful for predicting traits that are costly to phenotype when a correlated trait less expensive to phenotype was available^45^. In our study, when the prediction accuracy of the single-environment multivariate model MTM.UN was compared with the baseline model G-BLUP, several combinations of related and unrelated traits led to increased accuracy. For the related traits with a high phenotypic correlation (Figure 1a), prediction of traits measured at one location were often improved when a related trait measured across different locations was included. This was the case for the combination of floral emergence with full flowering and end of flowering and for single fruit weight combined with fruit diameter, fruit length, maximum fruit size and fruit volume. Inclusion of soluble solids content in MTM.UN resulted in increased prediction accuracy for russet cover, although the traits showed only a moderate correlation and no obvious explanation for this result could be found. Our study supports the potential of multivariate models to borrow information that correlated traits provide about one another and identified trait combinations that can be successful under the multivariate setup.

In place of the correlated traits, environments of a single trait can be implemented in a multivariate model^46^. The average prediction accuracy over all traits was ~0.04 lower in the multi-environment multivariate (MTM.FA) than in the multi-environment univariate genomic prediction models (G-BLUP.E and G-BLUP.E.G×E). Compared to G-BLUP.E and G-BLUP.E.G×E, the MTM.FA showed the potential to perform equally well for six (CV1) and three traits (CV2) and was able to outperform both models for two (CV1) and five traits (CV2). In cases where MTM.FA outperformed G-BLUP.E and G-BLUP.E.G×E, a very limited increase in prediction accuracy of 0.01 was found for all traits but trunk increment, for which the increase was equal to 0.07 under the second cross-validation scenario. Except for the noticeable increase in prediction accuracy for trunk increment that could not be explained by our analyses, the performance of MTM.FA was similar to G-BLUP.E and G-BLUP.E.G×E, which establishes the multivariate model as a useful tool for multi-environment genomic prediction in apple.

### Two approaches to genomic prediction addressed with cross-validation scenarios

The cross-validation scenarios CV1 and CV2 were applied with multi-environment genomic prediction models to test two genomic prediction approaches typically faced in breeding. The CV1 imitated evaluation of breeding material that was yet untested in field trials. The CV2 was implemented to simulate incomplete field trials where breeding material was evaluated in some but not all target environments. More specifically, the CV2 investigated a situation where the breeding material has been evaluated at one location (the breeding site, in this case Switzerland) and the material’s potential over other European sites was predicted without its assessment in a multi-environment trial, which may increase selection efficiency at latter stages of evaluation. As CV2 provided more phenotypic information to the models than CV1, a higher genomic prediction accuracy was found under CV2 when compared with CV1, which was anticipated^33,42^. The CV2 was tested by calibrating the model with Swiss observations only. The application of CV2 could be extended to other apple REFPOP locations to provide useful information for the breeding programs located at these sites. The choice of cross-validation scenario did not affect the general ranking of the average genomic prediction accuracies estimated for the evaluated traits.

### Implications for apple breeding

Phenotypic variance decomposition into genetic, environmental, G×E and residual effects was compared with the results of GWAS and genomic prediction as well as heritability estimates. The comprehensive comparison indicated three classes of traits with contrasting genetic architecture and prediction performance. Characteristics of these trait classes and proposals for their efficient prediction strategies are described in the following paragraphs.

The first class included harvest date and red over color that showed a few loci of large effect and some additional loci of low effect, the highest prediction accuracies, and the highest across-environment clonal-mean heritability among all traits. Both traits showed a very high proportion of the genotypic effect explaining ~75% of the phenotypic variance. For harvest date and red over color, the marker with the largest effect explained 52% and 59% of the phenotypic variance and all marker effects in genomic prediction captured together 88% and 85% of the phenotypic variance (i.e., genomic heritability of 0.88 and 0.85), respectively. Selection for these traits exhibiting a strong genetic effect of one locus could be done using marker-assisted selection, although only a part of the variance would be explained by a single marker. Better results can be achieved using genomic prediction, as this was able to explain a substantially larger amount of the phenotypic variance. Other traits such as fruit firmness, titratable acidity, end of flowering or traits related to fruit size and water core were grouped in the same cluster as harvest date and red over color (Figure 6). These traits showed a strong genotypic effect and a comparably low effect of environment and G×E, suggesting that selection for the traits would be efficient when performed using single-environment genomic prediction models rather than multi-environment prediction.

The second class of traits was represented by floral emergence and trunk diameter displaying a high proportion of the environmental effect (~70%) and a similar ratio of variance explained by genotypic effects compared to variance explained by G×E effects (~2.5). The genomic prediction accuracy did not considerably deviate from the average accuracy over all traits. Several marker associations with these traits were identified using location-specific GWAS. However, in the across-location GWAS, only one association explaining a very small part of phenotypic variance (floral emergence) or no association (trunk diameter) were discovered. Consequently, such traits predominantly driven by the effect of environment can be successfully selected based on genomic prediction, but the lack of associations stable across environments limits the applicability of marker-assisted selection to this class of traits.

In the third class, the productivity traits (flowering intensity, weight of fruits and number of fruits) showed the largest proportion of variance explained by G×E (~20%), with similar amounts of variance explained by genotypic effects for weight of fruits and number of fruits, but half as much variance explained by genotypic effects for flowering intensity (Figure 6). As a consequence, only flowering intensity showed higher prediction accuracy with G-BLUP.E.G×E than G-BLUP.E model. As shown above, the G×E should be considered when making predictions across environments for traits responding positively to the G-BLUP.E.G×E model, but G-BLUP.E may be sufficient for other traits to account for the environmental effects. To our knowledge, this is the first report of genomic prediction for apple yield components and our results can aid the establishment of productivity predictions in apple breeding. Other traits falling within the same cluster as the productivity traits, namely full flowering, ground color, yellow color, soluble solids content, trunk increment, and traits related to bitter pit and russet, showed a pronounced effect of environment and/or G×E (Figure 6). Multi-environment genomic prediction models can be efficient when applying genomic selection to these traits.

The decision to apply either marker-assisted or genomic selection can be based on genetic architecture of traits of interest and resources available in a breeding program. For breeding of yet genetically unexplored traits, variance decomposition of historical phenotypic data prior to genomic analyses may help describe trait architecture, assign traits to one of the three classes described in the previous paragraphs, and finally determine the most appropriate method of genomics-assisted breeding. From all traits explored in this study, the marker-trait associations with large and stable effects across environments found for harvest date, flowering intensity, green color, red over color, titratable acidity, fruit firmness and trunk increment could be implemented into DNA tests for marker-assisted selection. These tests would allow for a reduction of labor costs in a breeding program when removing inferior seedlings without phenotyping^7^. Although generally requiring more statistical competences than marker-assisted selection, genomic selection can make use of both large- and low-effect associations between markers and traits when accommodating thousands of marker effects in a single genomic prediction model. For all studied traits, our results showed that marker effects estimated in genomic prediction were able to capture a larger proportion of the phenotypic variance than individual markers associated with the traits. Therefore, genomic selection should become the preferred method of genomics-assisted breeding for all quantitative traits explored in this study to ultimately increase their breeding efficiency and genetic gain.

## Conclusion

This study laid the groundwork for marker-assisted and genomic selection across European environments for 30 quantitative apple traits. The apple REFPOP experimental design facilitated identification of a multitude of novel and known marker-trait associations. Our multi-environment trial provided accurate genomics-estimated breeding values for apple genotypes under various environmental conditions. Limited G×E detected in this work suggested consistent performance of genotypes across different European environments for most studied traits. Utilizing our dataset, more efficient selection of traits related to yield may lead to higher productivity and increased genetic gain in the future^37^. Improved fruit quality would appeal to consumers and tree phenology could be synchronized with current and future climates to secure production. The genomic prediction models developed here can be readily used for selecting germplasm in breeding programs, thus providing breeders with tools increasing selection efficiency. Beside the apple REFPOP, one other large multi-environment reference population for fruit trees, the PeachRefPop^64^, was designed in Europe. Application of our study design to other horticultural crops such as peach can promote broader use of genomics-assisted breeding in the future.

## Methods

### Plant material

The apple REFPOP was designed and established by the collaborators of the FruitBreedomics project^65^ as described by Jung et al.^36^. The apple REFPOP consists of 534 genotypes from two groups of diploid germplasm. The accession group consists of 269 accessions of European and non-European origin representing the diversity in cultivated apple. The progeny group of 265 genotypes stemmed from 27 parental combinations produced in the current European breeding programs. In 2016, the apple REFPOP was planted in six locations representing several biogeographical regions in Europe, in (i) Rillaar, Belgium, (ii) Angers, France, (iii) Laimburg, Italy, (iv) Skierniewice, Poland, (v) Lleida, Spain and (vi) Wädenswil, Switzerland (one location per country). Every genotype was replicated at least twice per location. All plants included in this study were treated with agricultural practice common to each location. Calcium spraying was avoided due to its influence on bitter pit. Flowers were not thinned, but the fruits were hand-thinned after the June fruit drop and up to two apples per fruit cluster were retained.

### Genotyping

A high-density genome-wide SNP marker dataset was produced as reported by Jung et al.^36^. Briefly, SNPs from two overlapping SNP arrays of different resolution, (i) the Illumina Infinium^®^ 20K SNP genotyping array^20^ and (ii) the Affymetrix Axiom^®^ Apple 480K SNP genotyping array^21^, were curated and then joined applying imputation with Beagle 4.0^66^ using the recently inferred pedigrees^4^. Non-polymorphic markers were removed to obtain a set of 303,148 biallelic SNPs. Positions of SNPs were based on the apple reference genome obtained from the doubled haploid GDDH13 (v1.1)^16^.

### Phenotyping

Thirty phenotypic traits related to phenology, productivity, fruit size, outer fruit, inner fruit, and vigor were evaluated at up to six locations of the apple REFPOP during up to three seasons (2018–020). Trunk diameter was measured in 2017 in some locations, enabling for a trunk increment calculation for 2018. The traits were recorded as described in the Supplementary Methods. Two phenology traits measured in 2018, i.e., floral emergence and harvest date, were previously analyzed by Jung et al.^36^.

### Phenotypic data analyses

Spatial heterogeneity was modeled separately for each trait and environment (nested factor of location and year) using the spatial analysis of field trials with splines (SpATS) to account for the replicate effects and differences due to soil characteristics^67^. Phenotypic values of traits adjusted for spatial heterogeneity within each environment were estimated at the level of trees (adjusted phenotypic values of each tree) and genotypes (adjusted phenotypic values of each genotype)^36^.

The general statistical model for the following phenotypic data analyses fitted via restricted maximum likelihood (R package lme4^68^) was:

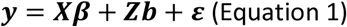

where ***y*** was a vector of trait phenotypes, ***X*** the design matrix for the fixed effects, ***β*** the vector of fixed effects, ***Z*** the design matrix for the random effects, ***b*** the vector of random effects and ***ε*** the vector of random errors. The ***b*** was a *q* × 1 vector assuming ***b*** ~ *N*(0, **Σ**) where **Σ** was a variance-covariance matrix of the random effects. The assumptions for the *N* × 1 vector of random errors were 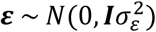 with *N* × *N* identity matrix ***I*** and the variance 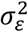, the *N* being the number of trees.

To assess the reliability of environment-specific data, a random-effects model was first fitted separately for each trait and environment to estimate an environment-specific clonal mean heritability. Applying the Equation 1, the response ***y*** was a vector of the raw (non-adjusted) phenotypic values of each tree. On the place of ***X***, a vector of ones was used to model the intercept *β*. The vector of genotypes acted as a random effect in ***Z***. The environment-specific clonal mean heritability was calculated from the variance components of the random-effects model as:

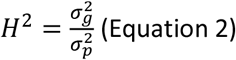

where the phenotypic variance 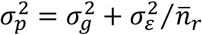 was obtained from the genotypic variance 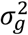, error variance 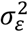 and the mean number of genotype replications 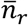. The environment-specific clonal mean heritability was used to eliminate location-year-trait combinations with a heritability value below 0.1. For the remaining location-year combinations, a mixed-effects model following the Equation 1 was fitted to the vector of the adjusted phenotypic values of each tree as response (***y***). The effects of environments, i.e., combination of location and years, were used as fixed effects and the effects of genotypes and genotype by environment interactions as random effects. Estimated variances of the model components were used to evaluate the across-environment clonal mean heritability calculated using the Equation 2 with the phenotypic variance estimated as:

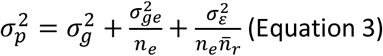

where 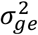 was the genotype by environment interaction variance and *n_e_* represented the number of environments.

An additional mixed-effects model following the Equation 1 was fitted to the adjusted phenotypic values of each tree (***y***) using the effects of location, year and their interaction as fixed effects and the effects of genotypes as random effects. Due to the skewness of their distributions, ***y***-values of the traits weight of fruits, number of fruits and trunk diameter were log-transformed. BLUPs 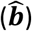 extracted from the model were further denoted as across-location BLUPs. To estimate the location-specific BLUPs, a model according to the Equation 1 was fitted with a subset of the adjusted phenotypic values of each tree from single locations (***y***) using the effects of years as fixed effects and the effects of genotypes as random effects. The across-location BLUPs and the adjusted phenotypic values of each genotype were used to assess phenotypic correlation as the Pearson correlation between pairs of traits and between pairs of environments within traits, respectively. The across-location BLUPs with the addition of location-specific BLUPs for traits measured at a single location were further denoted as the main BLUPs. In the main BLUPs, the missing values were replaced with the mean of the BLUPs of the same trait and the data were scaled and centered to finally estimate a principal component analysis biplot^69^, where multivariate normal distribution was assumed for the ellipses.

### Genome-wide association studies

The Bayesian-information and linkage-disequilibrium iteratively nested keyway (BLINK)^70^ implemented in the R package GAPIT 3.0^71^ was applied using the genomic matrix ***M***, an *n*×*m* matrix for a population of size *n* = 534 genotypes (i.e., accessions and progeny) with *m* = 303,148 markers, with across-location BLUPs (across-location GWAS) or location-specific BLUPs (location-specific GWAS) as the response. BLINK was used with two principal components and the minor allele frequency threshold was set to 0.05. Marker-trait associations were identified as significant for p-values falling below a Bonferroni-corrected significance threshold *α*\* = *α/m* with *α* = 0.05 (-*log*_10_(*p*) > 6.74). The proportion of phenotypic variance explained by each significantly associated SNP was assessed with a coefficient of determination (*R*^2^). The *R*^2^ was estimated from a linear regression model, which was fitted with a vector of SNP marker values (coded as 1, 2, 3) as predictor and either the across-location BLUPs or location-specific BLUPs as response. GWAS based on the across-location BLUPs with the addition of location-specific BLUPs, in cases where traits were measured at a single location only, was further denoted as the global GWAS. The position of the last SNP on a chromosome was used to estimate chromosome length, which was used to divide each chromosome into three equal segments, i.e., top, center and bottom. The marker-trait associations were assigned to these chromosome segments based on their positions.

Previous reports on QTL mapping and GWAS in apple were reviewed to perform an extensive comparison with our GWAS results (Supplementary Table 4). Published results for traits measured similarly to the traits studied in the present work were considered, with the traits being assembled into trait groups: harvest time (harvest date and similar), flowering time (floral emergence, full flowering, end of flowering and similar), productivity (flowering intensity, weight of fruits, number of fruits and similar), fruit size (single fruit weight, fruit diameter, fruit length, maximum fruit size, fruit volume and similar), ground color (ground color, yellow color and similar), over color (red over color, green color and similar), bitter pit (bitter pit frequency, bitter pit grade and similar), russet (russet cover, russet frequency overall, at stalk, on cheek and in the eye and similar), acidity (titratable acidity and similar), sugar (soluble solids content and similar), firmness (fruit firmness and similar), water core (water core frequency, water core grade and similar) and trunk (trunk diameter, trunk increment and similar). The positions of published associations within respective chromosomes were visually assigned to the three chromosome segments, i.e., top, center and bottom. The total number of markers used was recorded (Supplementary Table 4). Where the number of overlapping markers between the maternal and paternal linkage maps was not provided in a publication, the marker numbers for both maps were summed.

In the global GWAS results, the allele frequency was studied over generations. The ancestors of genotypes were identified making use of the apple pedigrees of Muranty et al.^4^. For all significant marker-trait associations from the global GWAS, frequency of the allele associated with increased phenotypic value was estimated for the progeny group and for its five ancestor generations. To represent the ancestors, the allele frequency was estimated for the 30 accessions of them included in the apple REFPOP. For major significant marker-trait associations with *R*^2^ > 0.1 reported in the global GWAS, linkage disequilibrium was estimated as squared Pearson’s correlations in a window of 3,000 markers surrounding each of the associations. A smaller window size was used for associations located towards the end of a chromosome.

A mixed-effects model following the Equation 1 was fitted to the vector of the adjusted phenotypic values of each tree as response (***y***) using the effects of environments as fixed effects and the effects of genotypes, genotype by environment interactions, and additional effects for each SNP significantly associated with the trait (a factor of the respective SNP values in ***M***) as random effects. In cases where traits with no marker-trait associations were found in the global GWAS, the additional random effects of significantly associated SNPs were omitted from the model. The mixed-effects model for every trait was used to estimate proportions of phenotypic variance explained by the model components as described in Jung et al.^36^. The proportions of phenotypic variance explained by the random effects of genotypes and significantly associated SNPs were summed to obtain the proportion of variance explained by a genotypic effect. The proportions of phenotypic variance explained by genotypic, environmental, genotype by environment interaction, and residual effects were scaled and centered to be finally used for discovering similarities between the traits. For this purpose, a hierarchical clustering following Ward^72^ was applied to the distance matrix of the set of effects. The number of clusters was estimated from a dendrogram, which was cut where the distance between splits was the largest.

### Genomic prediction

The general statistical model for genomic prediction was

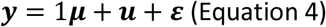

where ***y*** was a vector of trait phenotypes, ***μ*** was an intercept, ***u*** represented a vector of random effects and ***ε*** was a vector of residuals. Different vectors of ***y*** and assumptions for ***u*** and ***ε*** were used across eight single- and multi-environment genomic prediction models.

#### Single-environment genomic prediction

The single-environment genomic prediction models were fitted after the environmental effects were accounted for during the phenotypic data analysis, a process also called two-step genomic prediction. Therefore, the across-location BLUPs and location-specific BLUPs acted here as phenotypes from a single environment. Four univariate prediction models and one multivariate model were implemented. First, regression with random forest (RF) was performed^73^. In this and the following three univariate models, the response ***y*** was defined as a *n* × 1 vector of the main BLUPs. The centered and scaled additive genomic matrix ***M***, an *n*×*m* matrix for a population of size *n* = 534 with *m* = 303,148 markers, was used as further input. The number of trees to grow in the RF was 500 and the number of variables randomly sampled as candidates at each split was (rounded down) *mtry* = *m*/3. Second, BayesCπ was applied^74^, where the random marker effects 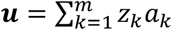 with *z_k_* an *n* × 1 vector of the number of copies of one allele at the marker *k* and *a_k_* being the additive effect of the marker *k*. The prior for *a_k_* depended on the variance 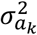 and the prior probability *π* that a marker *k* had zero effect, the priors of all marker effects having a common variance 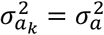. The *π* parameter was treated as an unknown with uniform(0,1) prior. The random vector of residual effects followed a normal distribution 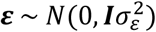 with *n*×*n* identity matrix ***I*** and the variance 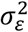. Third, the Bayesian reproducing kernel Hilbert spaces regression (RKHS) was implemented using a multi-kernel approach^75^. The multi-kernel model was fitted with *L* = 3 random marker effects 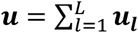 following a distribution 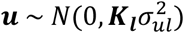, with ***K_l_*** being the reproducing kernel evaluated at the *l*th value of the bandwidth parameter *h* = {*h*_1_,…, *h_L_*} = {0.1,0.5,2.5} and the variance 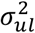. For each random effect, the kernel matrix ***K*** = {*K*(*x_i_, x_i′_*)} was an *n*×*n* matrix *K*(*x_i_,x_i′_*) = exp{-*h*×*D_ii′_*}, where 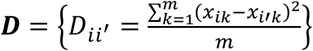 was the average squared-Euclidean distance matrix between genotypes, and *x_ik_* the element on line *i* (genotype *i*) and column *k* (*k*th marker) of the centered and scaled additive genomic matrix ***M***. The residual effect assumed 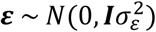. Fourth, from the centered and scaled additive genomic matrix ***M***, the genomic relationship matrix ***G*** was computed as ***G*** = ***MM’**/m* and used to fit the genomic-BLUP (G-BLUP) model applying a semi-parametric RKHS algorithm, with the random marker effects following 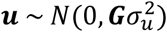 with variance 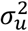 and the model residuals assuming 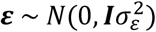^76^. Fifth, a multivariate model with an unstructured covariance matrix of the random marker effect (here abbreviated as MTM.UN) was fitted for chosen pairs of traits using the Bayesian multivariate Gaussian model environment MTM (http://quantgen.github.io/MTM/vignette.html). The main BLUPs acted as the response ***y***, which was a vector of length *n* · *t* with *t* = 2 being the number of traits used in the model. The vector of the random marker effects followed ***u*** ~ *N*(0, ***U*** ⊗ ***G***) where ***U*** was an unstructured covariance matrix of the random marker effect with dimension *t* × *t*. Model residuals assumed ***ε*** ~ *N*(0, ***R*** ⊗ ***I***) with *R* being an unstructured covariance matrix of the residual effect. To choose the pairs of traits for MTM.UN, a G-BLUP model was applied using all genotypes to estimate genomic BLUPs, which were then used to obtain pairwise genomic correlations between traits. The pairs with the genomic correlations larger than 0.3 were retained for the MTM.UN analysis. In case a trait was included in more than one pair of traits, the result for the pair with the highest average predictive ability for this trait was reported.

BayesCπ, RKHS, G-BLUP and MTM.UN were applied with 12,000 iterations of the Gibbs sampler, a thinning of 5, and a burn-in of 2,000 discarded samples. With all models, a five-fold cross-validation repeated five times was performed, generating 25 estimates of prediction accuracy. The folds were chosen randomly without replacement to mask phenotypes of 20% of the genotypes in each run. Prediction accuracy was estimated as a Pearson correlation coefficient between phenotypes of the masked genotypes and the predicted values for the same genotypes. The RF model was implemented in the R package ranger^77^, the models BayesCπ, RKHS and G-BLUP in the R package BGLR^78^ and the MTM.UN model in the R package MTM (http://quantgen.github.io/MTM/vignette.html).

#### Multi-environment genomic prediction

Two univariate multi-environment genomic prediction algorithms and one multivariate multi-environment algorithm were implemented, the response ***y*** being a vector of the adjusted phenotypic values of each genotype of length *n* × *r* with *r* equal to the number of environments (nested factor of location and year). The two univariate multi-environment models reported by Lopez-Cruz et al.^42^ and implemented in the R package BGLR^78^ were applied to explore the effects of genotypes, environments and their interaction in genomic prediction. Of the two models, the across-environment G-BLUP model (G-BLUP.E) assumed that marker effects were constant across environments. The random marker effects followed 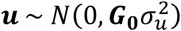 where ***G***_0_ = ***J*** ⊗ ***G***, the ***J*** being an *r* × *r* matrix of ones. The model residuals assumed 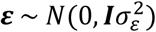. Additionally to the constant effects of markers across environments as assumed in the previous model, the marker by environment interaction G-BLUP model (G-BLUP.E.G×E) allowed the marker effects to change across environments, i.e., to borrow information across environments. The random marker effects were defined as ***u*** = ***u*_0_** + ***u*_1_** where 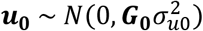 and ***u*_1_** ~ *N*(0, ***G*_1_**) with

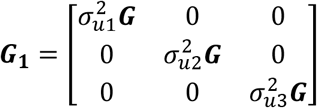

assuming *r* = 3 here for easier notation. The model residuals assumed 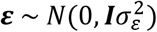. Finally, a multivariate multi-environment factor-analytic model (here abbreviated as MTM.FA) using the Bayesian multivariate Gaussian model environment implemented in the R package MTM (http://quantgen.github.io/MTM/vignette.html) was fitted to the data. As in the previous two models, phenotypes of the same trait from multiple environments acted as response, although this model was originally designed to analyze multiple traits. The traits measured at only one location during two seasons (full flowering, end of flowering, fruit volume, water core frequency and water core grade) were not modeled using MTM.FA because the analysis required at least three environments. The vector of the random marker effects assumed ***u*** ~ *N*(0, ***C*** ⊗ ***G***) where *C* was an *r* × *r* covariance matrix. For the factor analysis, the ***C*** = ***BB’*** + **Ψ** where ***B*** was a matrix of loadings (regressions of the original random effects into common factors) and **Ψ** was a diagonal matrix whose entries gave the variances of environment-specific factors. The loadings were estimated for all environments and the variance of the Gaussian prior assigned to the unknown loadings was set to 100. The model residuals assumed ***ε*** ~ *N*(0, ***R*** ⊗ ***I***) with ***R*** being an unstructured covariance matrix of the residual effect.

All three multi-environment genomic prediction models were applied with 12,000 iterations of the Gibbs sampler, a thinning of 5 and a burn-in of 2,000 discarded samples. The folds of a five-fold cross-validation were chosen randomly without replacement. The cross-validation was repeated under two scenarios. In the first cross-validation scenario (CV1), the phenotypes of 20% of the genotypes were masked across all environments. For the second cross-validation scenario (CV2), the phenotypes of 20% of the genotypes were masked across all environments except for three Swiss environments, i.e., phenotypes of all genotypes from the environments “CHE.2018”, “CHE.2019” and “CHE.2020” were used for model training. Ten traits were measured in only one location and therefore excluded from CV2 (i.e., full flowering, end of flowering, fruit diameter, fruit length, maximum fruit size, fruit volume, yellow color, green color, water core frequency and water core grade). Prediction accuracy was estimated as a Pearson correlation coefficient between the phenotypes of the masked genotypes and the predicted values for these genotypes. The correlations were estimated for each predicted environment separately.

### Genomic heritability

The BayesCπ model was applied for each trait as described before but trained with a full set of the main BLUPs as response. The genomic heritability *h*^2^ = *V_g_*/(*V_g_* + *V_e_*) was estimated as the proportion of phenotypic variance explained by the markers, where *V_g_* and *V_e_* represented the amount of phenotypic variance explained and unexplained by the markers, respectively^79,80^. The genomic heritability was calculated from the marker effects saved in each iteration and averaged over iterations to obtain the mean genomic heritability per trait.

## Data availability

All SNP genotypic data generated with the 480K array used in this study have been deposited in the INRAe dataset archive (https://data.inrae.fr/) at https://doi.org/10.15454/IOPGYF. All SNP genotypic data generated using the 20K array used in this study have been deposited in the INRAe dataset archive at https://doi.org/10.15454/1ERHGX. The raw phenotypic data generated during the study are available in the INRAe dataset archive at (TBA upon acceptance).

## Acknowledgements

The authors thank the field technicians and staff, especially Sylvain Hanteville, at INRAe IRHS and Experimental Unit (UE Horti), Angers, France, and technical staff at other apple REFPOP sites for the maintenance of the orchards and phenotypic data collection. We thank Dr. Graham Dow for English language editing. Phenotypic data collection was partially supported by the Horizon 2020 Framework Program of the European Union under grant agreement No 817970 (project INVITE: “Innovations in plant variety testing in Europe to foster the introduction of new varieties better adapted to varying biotic and abiotic conditions and to more sustainable crop management practices”). This work was partially supported by the project RIS3CAT (COTPA-FRUIT3CAT) financed by the European Regional Development Fund through the FEDER frame of Catalonia 2014-2020 and by the CERCA Program from Generalitat de Catalunya. We acknowledge financial support from the Spanish Ministry of Economy and Competitiveness through the “Severo Ochoa Programme for Centres of Excellence in R&D” 2016-2019 (SEV-20150533) and 2020-2023 (CEX2019-000902-S). C.D. was supported by “DON CARLOS ANTONIO LOPEZ” Abroad Postgraduate Scholarship Program, BECAL-Paraguay. We dedicate this paper to Prof. Edward Zurawicz of the National Institute of Horticultural Research in Skierniewice, Poland who co-promoted this study, but sadly recently passed away.

