## Supplementary Figures and Methods for "Genetic architecture and genomic prediction accuracy of apple quantitative traits across environments"

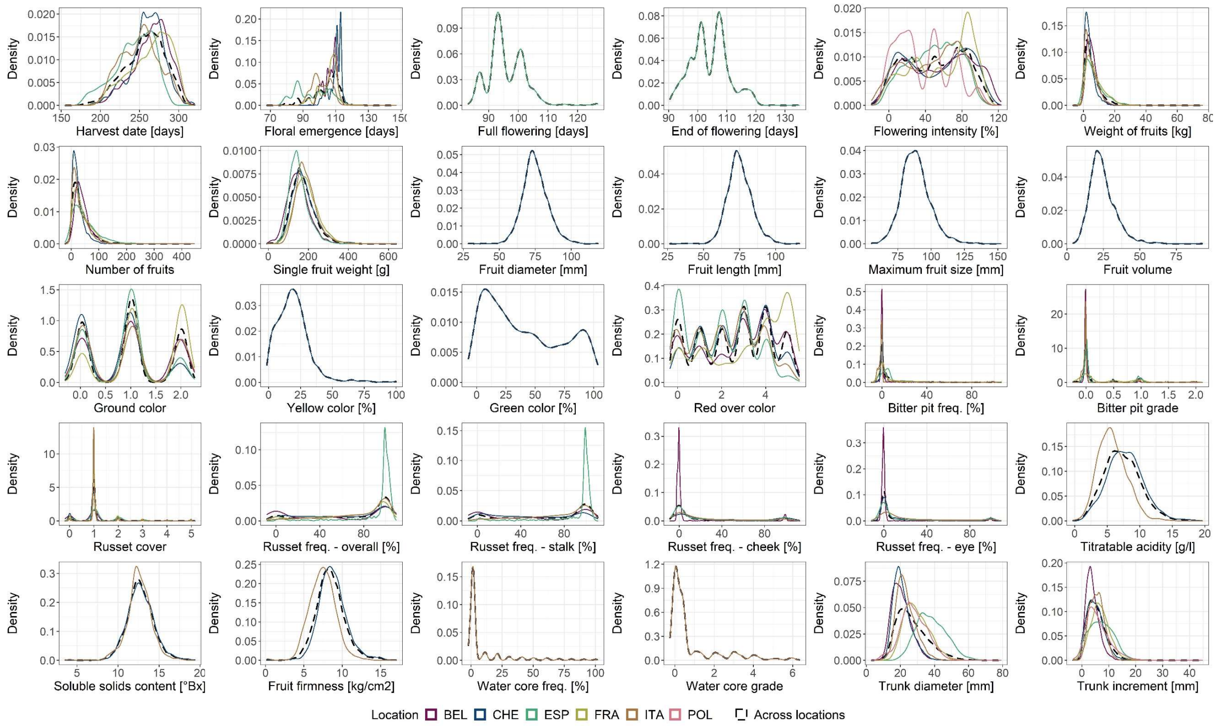


**Supplementary Figure 1:** Distributions of phenotypic values of traits adjusted for spatial heterogeneity within environments (adjusted phenotypic values of each tree), plotted per location.

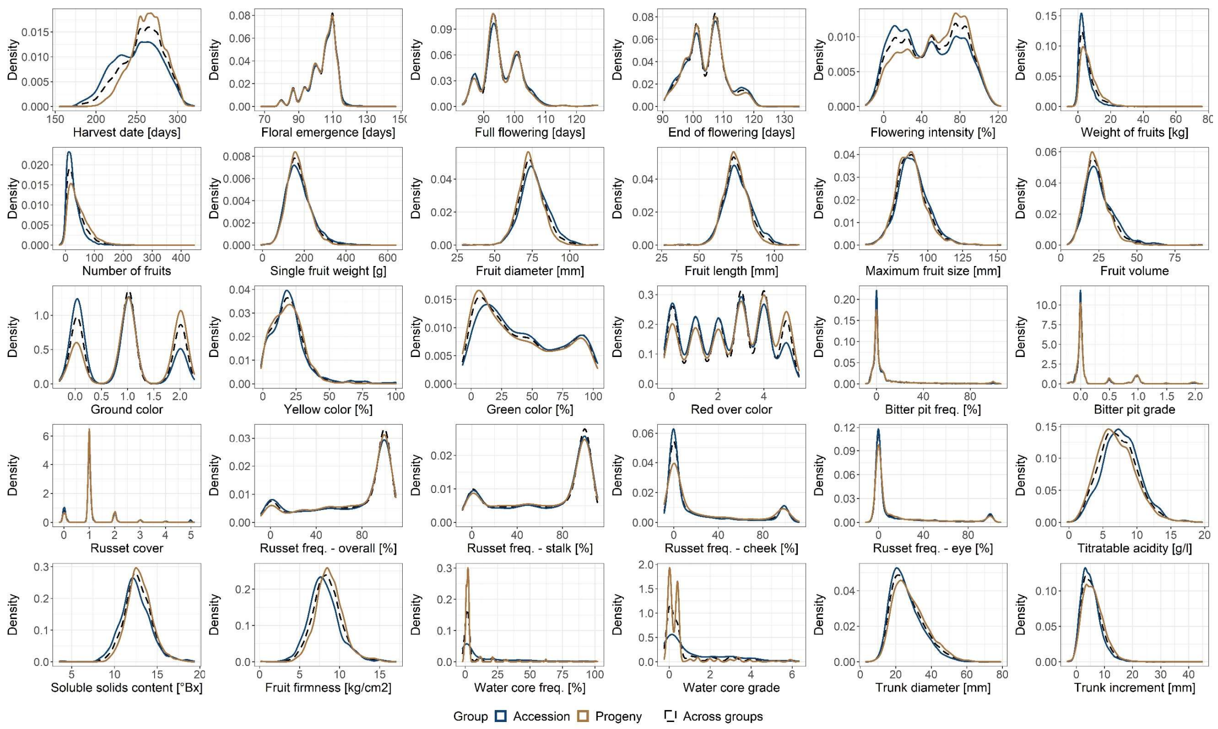


**Supplementary Figure 2:** Distributions of phenotypic values of traits adjusted for spatial heterogeneity within environments (adjusted phenotypic values of each tree), plotted per apple REFPOP group.

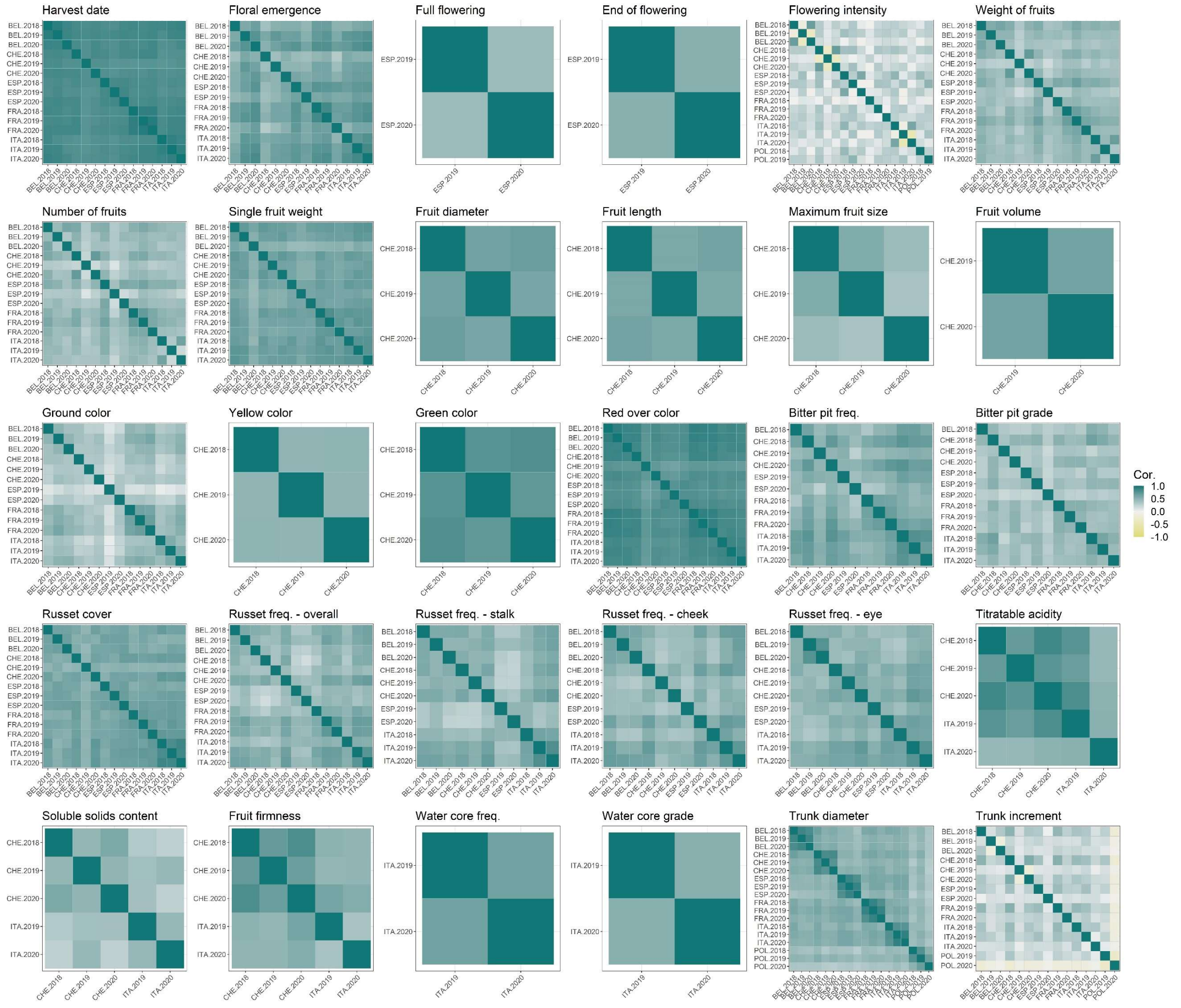


**Supplementary Figure 3:** Pairwise correlations of the adjusted phenotypic values of each genotype measured in different environments for individual traits.

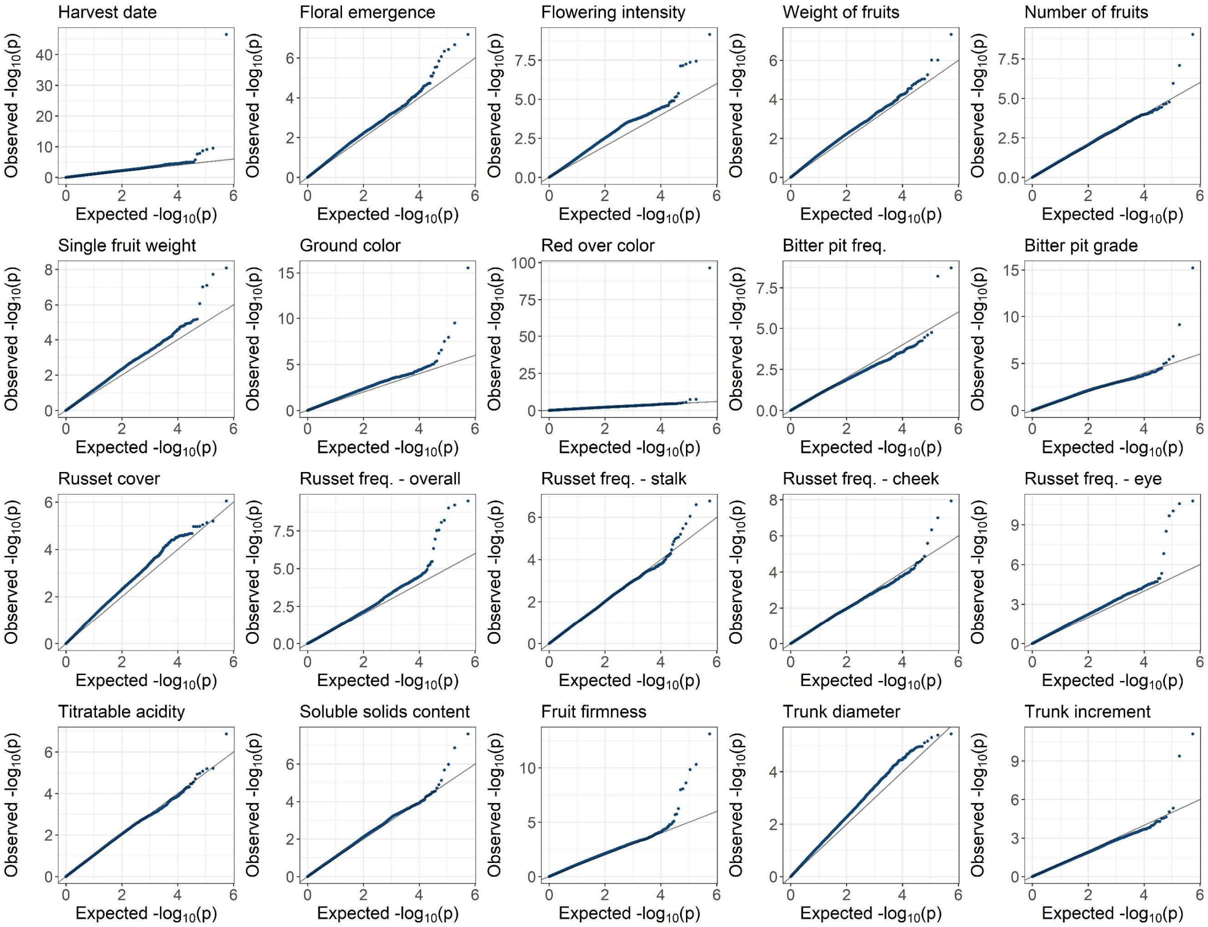


**Supplementary Figure 4:** QQ plots of the observed versus expected p-values for individual traits from the across-location GWAS.

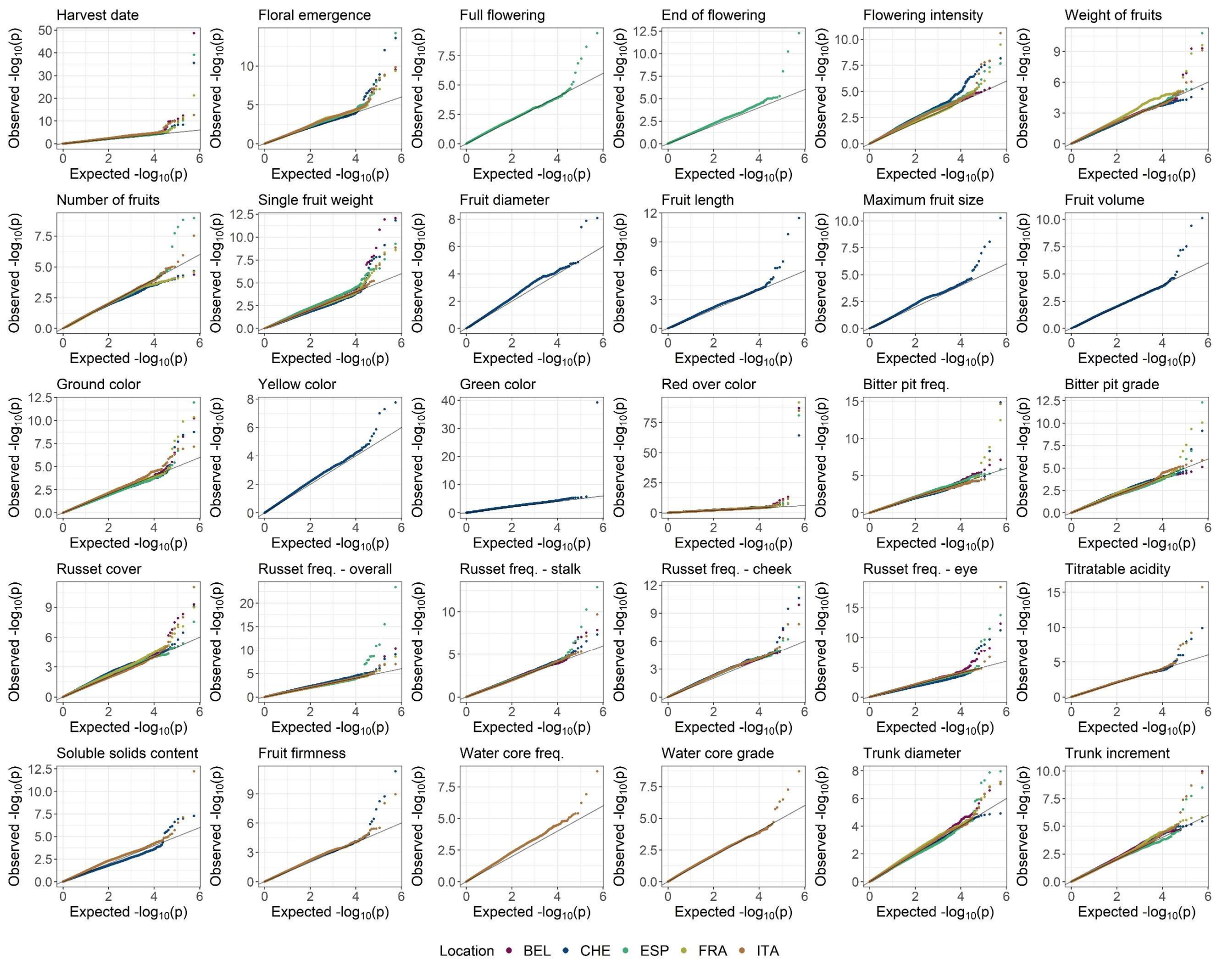


**Supplementary Figure 5:** QQ plots of the observed versus expected p-values from the location-specific GWAS for individual traits and locations.

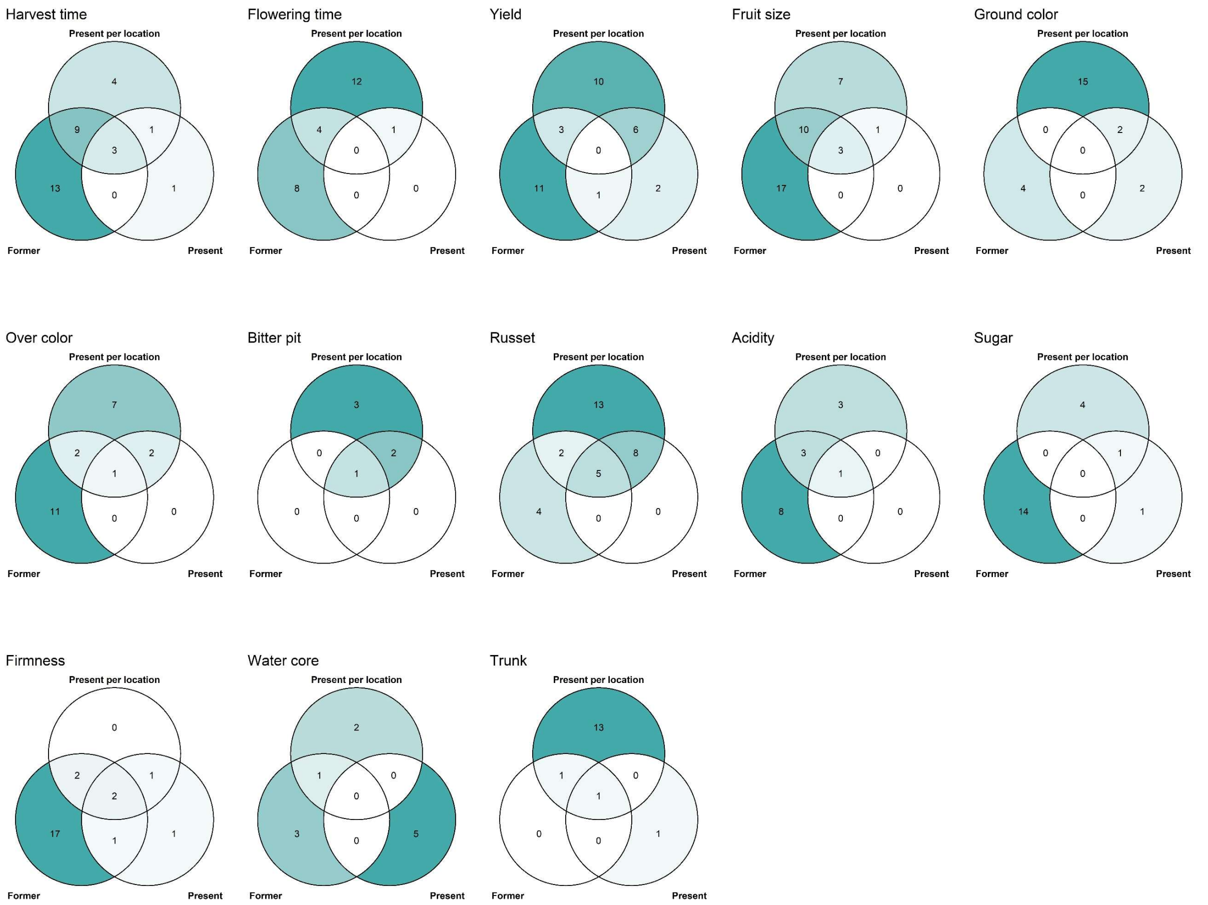


**Supplementary Figure 6:** Venn diagrams for each trait comparing the number of published associations (former, see also Supplementary Table 4) with the significant marker-trait associations from the across-locations GWAS (present, see also Supplementary Table 3) and location-specific GWAS (present (per location), see also Supplementary Table 3). The traits were assembled into trait groups based on their similarity. Color intensity reflects the number of associations per diagram area. The associations were assigned to chromosome segments (top, center, and bottom of a chromosome).

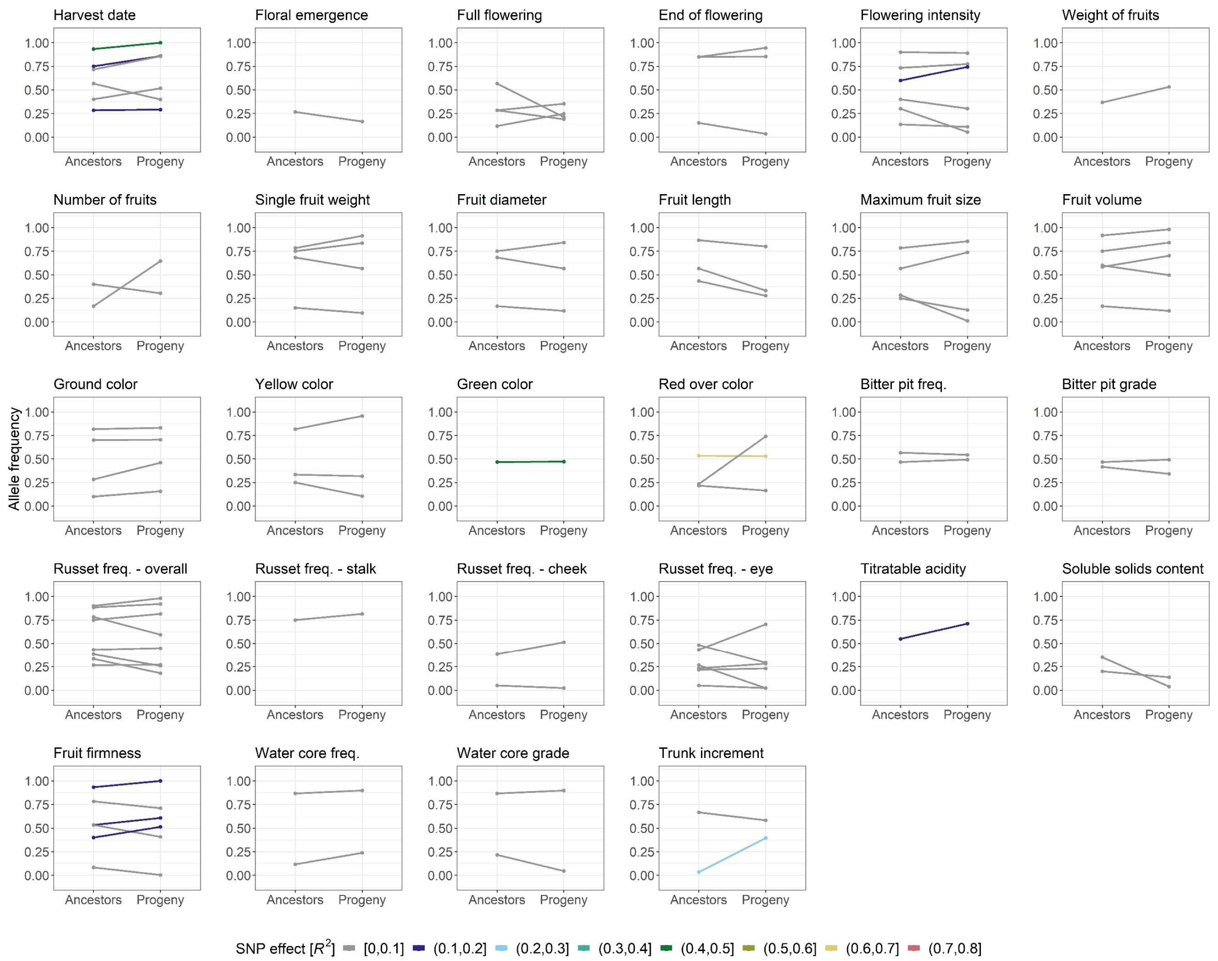


**Supplementary Figure 7:** Frequencies of alleles associated with increased phenotypic value for all significant marker-trait associations from the global GWAS. For the apple REFPOP progeny group (progeny) and its five ancestor generations (ancestors), the allele frequencies are shown as points connected with a line. Out of all known ancestors, the allele frequency was estimated for 30 accessions included in the apple REFPOP. Colors of the regression lines correspond to the part of phenotypic variance (*$R^2$*) explained by the associated SNPs.

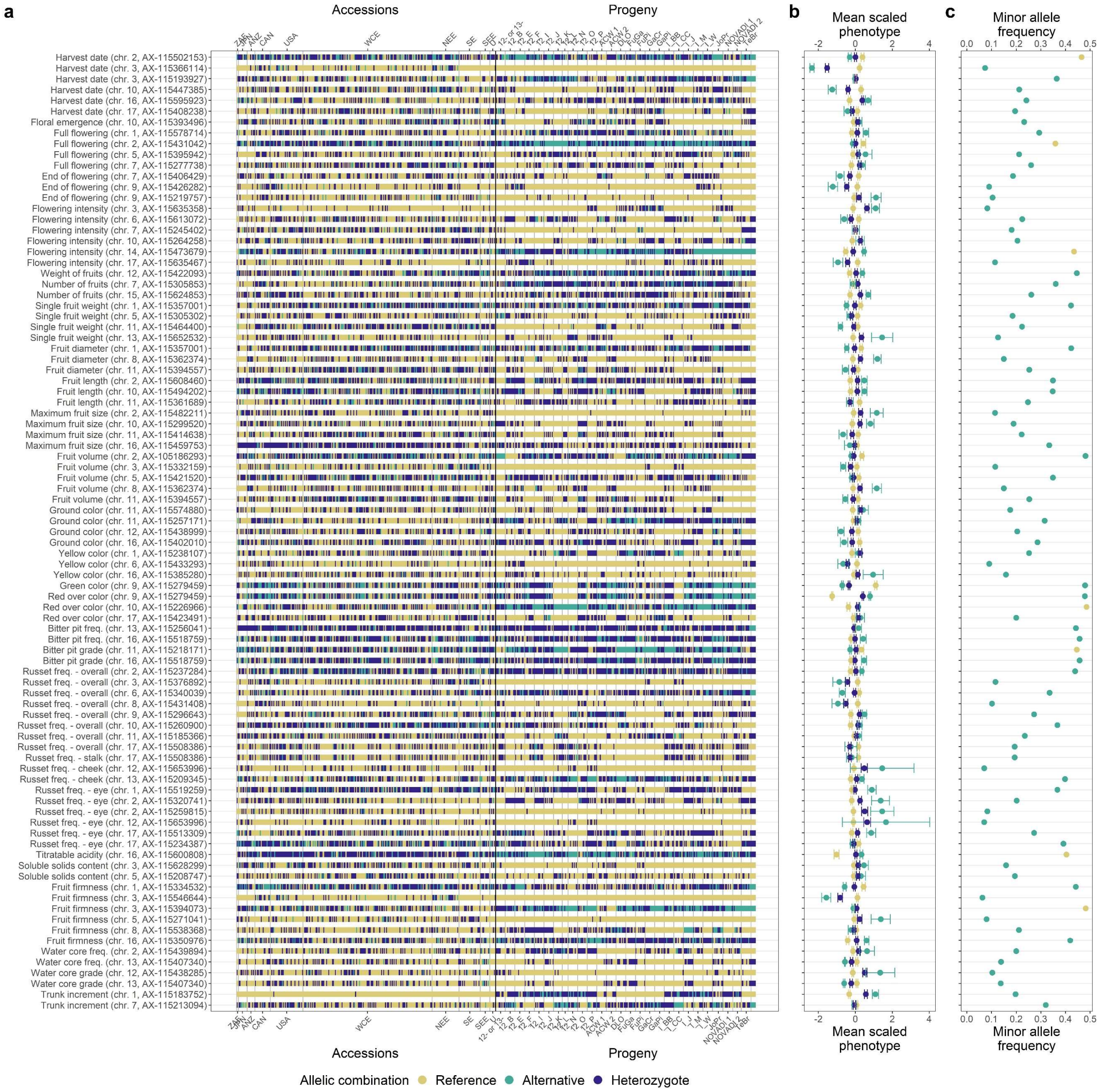


**Supplementary Figure 8: a** Allelic combinations carried by the apple REFPOP genotypes, sorted according to geographic origin of accessions (269) and affiliation of progeny (265) to parental combinations (the x-axis was labeled according to Supplementary Table 1 and 2 in Jung et al. 2020). **b** Mean scaled and centered phenotypic BLUPs of traits and their standard error for each allelic combination. **c** Frequency of the minor allele in the whole apple REFPOP. **a-c** The legend and y-axis are shared between plots. In c, the color of an allelic combination corresponds to an allele of the same name. Presented are associations from the global GWAS (across-location GWAS with the addition of location-specific GWAS for traits measured at a single location only).

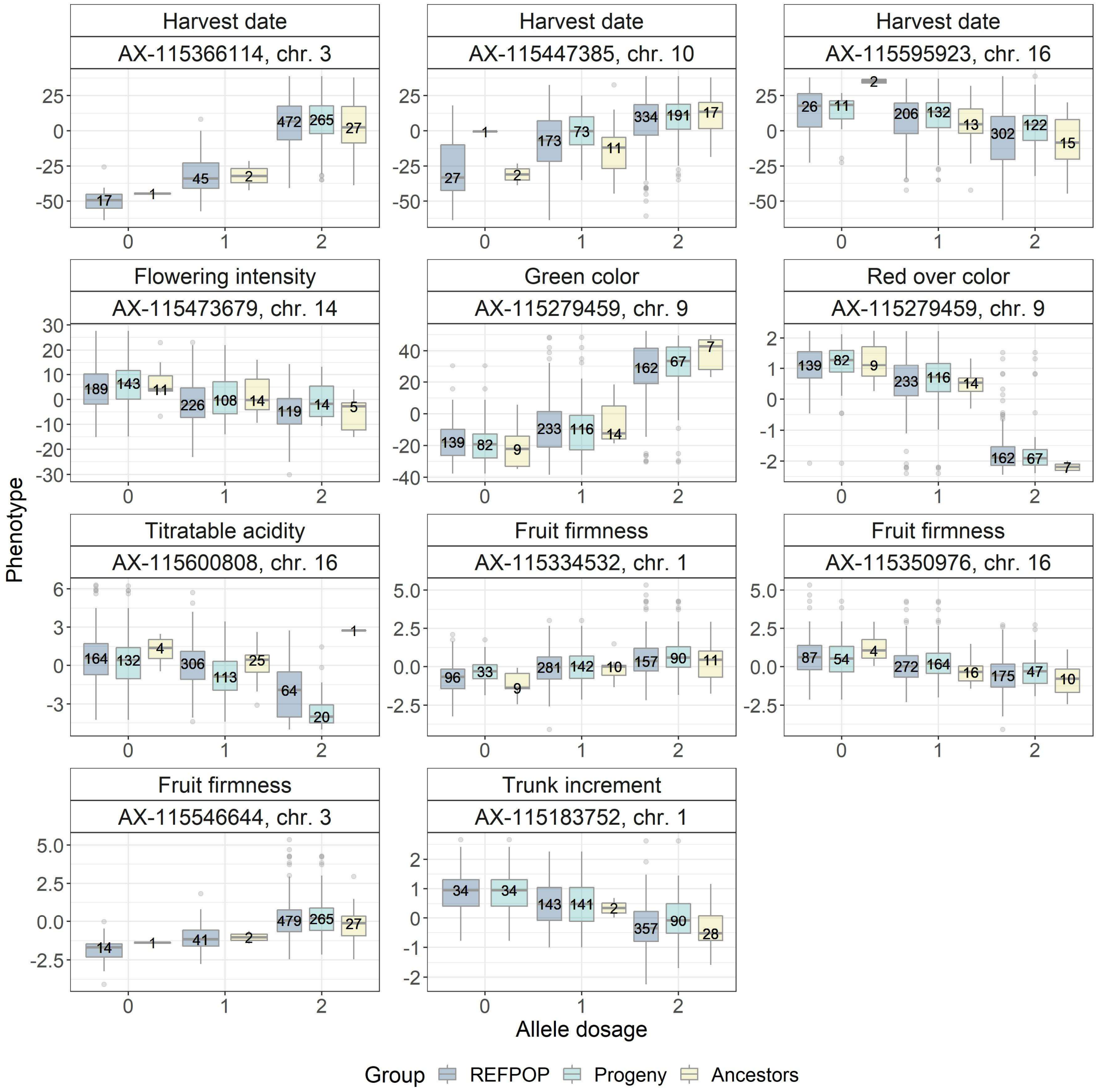


**Supplementary Figure 9:** Boxplots of phenotypes (across-location BLUPs) against dosage of the reference allele (0 – reference allele, 1 – heterozygote, 2 – alternative allele) for the major significant marker-trait associations, plotted for all apple REFPOP genotypes (REFPOP), the apple REFPOP progeny group (Progeny) and 30 ancestral accessions of the progeny group included in the apple REFPOP (Ancestors). Number of genotypes of a subgroup is shown in each box. Less than nine boxplots per trait indicate that not all specific allelic combinations were present in the respective group.

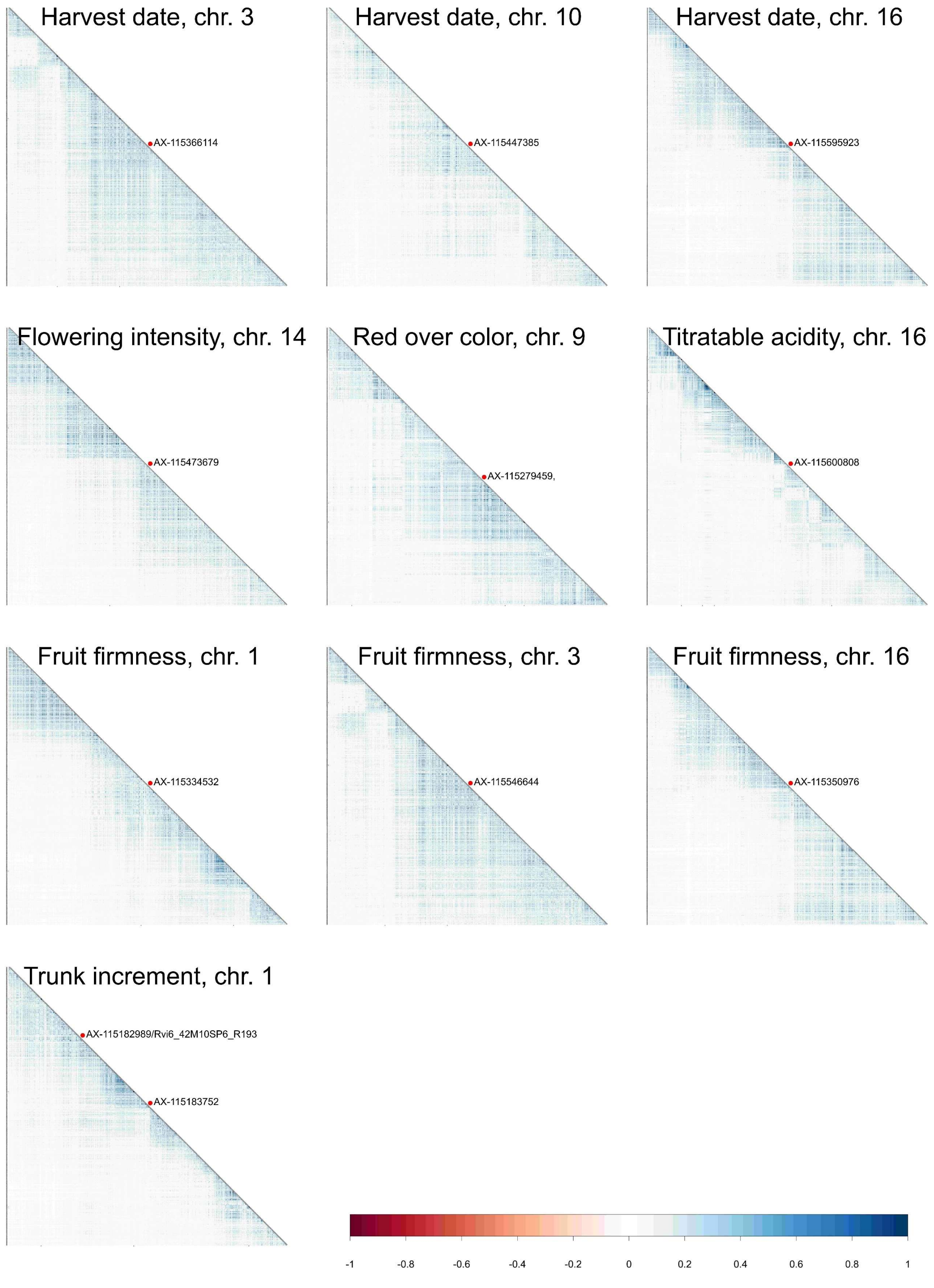


**Supplementary Figure 10:** Linkage disequilibrium estimated as squared Pearson's correlations in a window of 3,000 markers surrounding each of the major significant marker-trait associations. For the association with red over color, which corresponds to green color, only 2,736 markers were used due to the position of the association towards the end of chromosome nine. Position of a marker associated with apple scab resistance (*Rvi6*) is additionally shown in the plot of trunk increment. Physical size of the marker windows ranged between 4.1 Mb (harvest date, chromosome 10) and 5.8 Mb (trunk increment, chromosome 1).

### Supplementary methods

All traits were measured at the level of individual trees (genotype replicates). From the traits, 20 were measured at more than one location and ten were measured at one location only (see Supplementary Table 1 for trait-location combinations). Unless stated otherwise, the measurement of traits was performed on the whole crop (further denoted as produced fruits). The fruits fallen to the ground due to, e.g., over-ripeness or strong wind, were also collected unless they were heavily decayed or found too far away from the tree. Fruits that were not fully developed and of a very small size due to, e.g., aphids or second flowering, were discarded and not counted as produced fruits. Weight of fruits and outer fruit traits were scored on a fruit batch consisting of 20 fruits selected randomly or all produced fruits for trees bearing at least five and up to 20 fruits. Trees producing less than five fruits were not scored for traits measured on a fruit batch.

#### Phenology traits

*Floral emergence* was estimated as the date when the first 10% of flowers opened. *Full flowering* was estimated as the date when at least 50% of flowers were open. *End of flowering* was the date when all flower petals fell off. On *harvest date*, more than 50% of the produced fruits were physiologically fully mature. The ripeness of fruits for the estimation of the harvest date was determined by expert knowledge. Every date was converted to the number of days since the beginning of year of the measurement.

#### Productivity traits

*Flowering intensity* was the percentage of existing flowers from the maximum possible number of flowers. A tree would show the maximum possible number of flowers if flowering at each leafy shoot. The used scale was: grade 1 corresponded to 0%, grade 3 to 25%, grade 5 to 50%, grade 7 to 75% and grade 9 to 100%. To measure the *number of fruits*, all produced fruits were counted on harvest date. *Weight of fruits* in kilograms was measured with scales or a sorting machine (Greefa iQS4 v.1.0) from the full set of produced fruits or from the fruit batch. In case the weight of a fruit batch was measured, the weight of all fruits was calculated using the average fruit weight estimated from the batch multiplied by the number of all produced fruits.

#### Fruit size

*Single fruit weight* in grams was obtained by dividing the weight of all fruits in grams by the number of fruits. *Fruit diameter*, *fruit length*, *maximum fruit size* and *fruit volume* were estimated with the sorting machine for each produced fruit. To measure fruit diameter in mm, the sorting machine made twelve images per fruit, estimated the diameter from each image (along the horizontal axis), excluded the lowest and the largest value and averaged the remaining ten estimates. Fruit length in mm was obtained similarly to fruit diameter but using the vertical axis instead of the horizontal axis. Maximum fruit size in mm was derived from the fruit diameter estimates obtained from twelve images as an average of the second and the third largest values. Fruit volume was estimated using a ready-made algorithm of the sorting machine. Fruit diameter, fruit length and fruit volume were averaged across all produced fruits. For the maximum fruit size, the maximum value among all produced fruits was used.

**Outer fruit**

*Ground color* was visually estimated for a fruit batch assigning a grade between zero and two (Figure SM 1). *Red over color* represented a percentage of red fruit skin estimated visually. The assessment of red over color was performed on a fruit batch, where a grade between zero and five was assigned to the whole batch (Figure SM 2). *Yellow color* and *green color* were estimated automatically using the sorting machine as an average percentage of these colors on the skin of produced fruits.

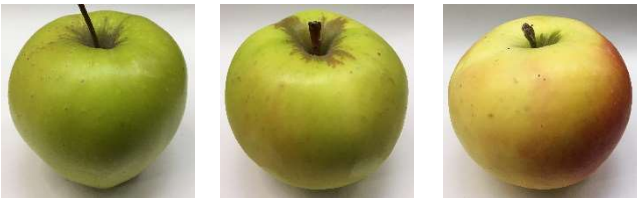

| green | green-yellow<br>yellow-green | yellow |
|---|---|---|
| (0) | (1) | (2) |

**Figure SM 1:** Scale for the assessment of ground color with the corresponding grades in brackets

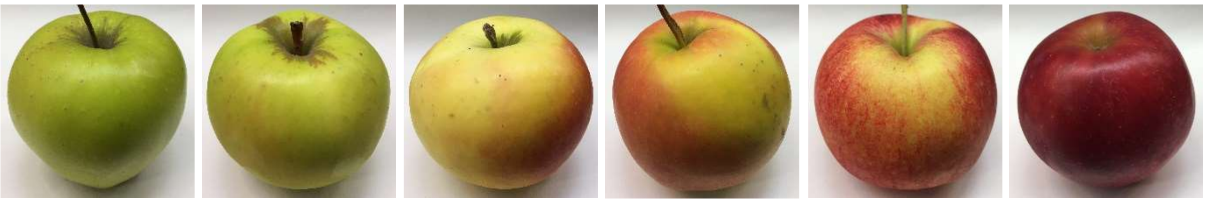

| absent | 1–20% | 21–40% | 41–60% | 61–80% | 81–100% |
|---|---|---|---|---|---|
| (0) | (1) | (2) | (3) | (4) | (5) |

**Figure SM 2:** Scale for the assessment of red over color with the corresponding grades in brackets

Bitter pit is a disorder characterized by discrete pits in the fruit flesh, which turn brown and desiccate over time[1]. *Bitter pit frequency* was measured as the number of apples in a batch that were showing symptoms of bitter pit. The frequencies were transformed into percentages, which represented proportions of the batch size. *Bitter pit grade* was determined for a fruit batch assigning a grade between zero and two (Figure SM 3).

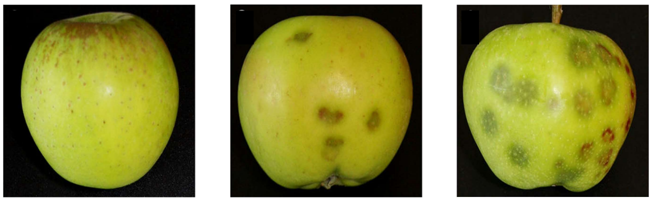

no symptoms (0) | medium symptoms (1) | very high symptoms (2)

**Figure SM 3:** Scale for the assessment of bitter pit grade with the corresponding grades in brackets, the scale was adapted from Buti et al.[1]

Russet is a fruit characteristic caused by small cracks in fruit skin, which develop during fruit growth and are replaced by cork tissue leading to rough brown skin. *Russet cover* was estimated as a percentage of fruit surface covered by the russet, estimated for a fruit batch assigning a grade between zero and five (Figure SM 4). A small patch of russet deep in the stalk cavity, which reached up to 50% of the depth of the cavity and in case of no other signs of russet on the fruit skin was considered an absent russeting (grade 0). *Overall russet frequency* was measured as the number of apples in a batch that were showing russet. *Russet frequency in the stalk basin*, *russet frequency on the cheek* and *russet frequency in the eye basin* were equal to the number of apples in a batch showing russet in the given area. All frequencies were transformed into percentages as proportions of the batch size.

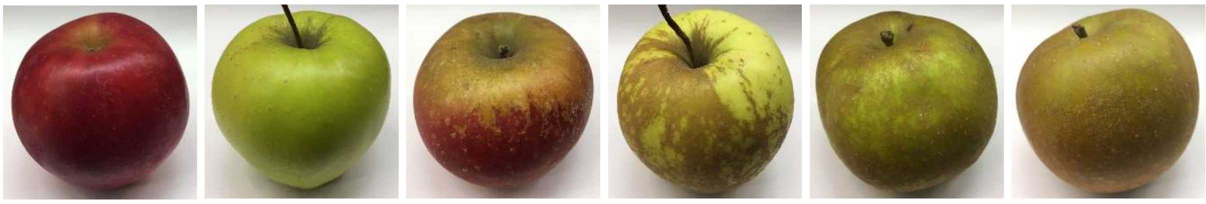

absent (0) | 1–20% (1) | 21–40% (2) | 41–60% (3) | 61–80% (4) | 81–100% (5)

**Figure SM 4:** Scale for the assessment of russet cover with the corresponding grades in brackets

**Inner fruit**

A sample of ten fruits was analyzed with an automated instrument Pimprenelle (Setop, France) within one week after the harvest date. A tree was omitted from the analysis with Pimprenelle in case it produced less than ten fruits (or less than five fruits in 2018 due to young age of trees linked with low production). *Fruit firmness* in g/cm$^2$ was measured on individual fruits of the sample using a penetrometer. Each fruit was then pressed to extract the juice, which passed onto a refractometer to measure *soluble solids content* as a refraction index in degrees Brix. Mean values of fruit firmness and soluble solids content were calculated per fruit sample, the refraction index of the first measured fruit was deleted by the Pimprenelle. All juice samples of the same fruit sample were pooled for the analysis of *titratable acidity*, which was measured in grams of titratable acid per liter of solution.

Water core is a disorder that is characterized by areas of fruit flesh soaked with water appearing as translucent tissue. *Water core frequency* was estimated as the number of apples in a sample of ten fruits that were showing symptoms of water core. *Water core grade* was visually scored from the sample of ten fruits assigning a grade between zero and six, the zero grade being equal to absence of water core in the sample (Figure SM 5).

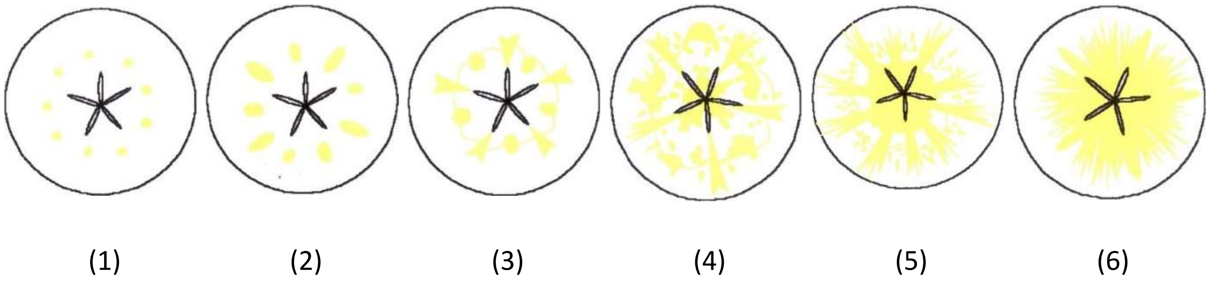


**Figure SM 5:** Scale for the assessment of water core grade with the corresponding grades in brackets (developed and applied at the Research Centre Laimburg, South Tyrol, Italy)

#### Vigor

Before the onset of flowering, *trunk diameter* in mm was measured 20 cm above the grafting point of trees using a digital caliper. Alternatively, trunk circumference was measured at the Italian site and transformed using the relation of circle's circumference to the diameter. *Trunk increment* in mm was obtained as difference between trunk diameter of the current and previous year.
